# Protrusion growth driven by myosin-generated force

**DOI:** 10.1101/2022.05.06.490961

**Authors:** Gillian N. Fitz, Meredith L. Weck, Caroline Bodnya, Olivia L. Perkins, Matthew J. Tyska

**Affiliations:** Department of Cell and Developmental Biology Vanderbilt University School of Medicine Nashville, TN 27232

**Keywords:** plasma membrane, actin, filopodia, cytoskeleton, microvilli, stereocilia, motor

## Abstract

Actin-based protrusions are found on the surface of all eukaryotic cells, where they support diverse biological activities essential for life. Models of protrusion growth hypothesize that actin filament assembly provides the mechanical force for bending the plasma membrane outward. However, membrane-associated myosin motors are also abundant in protrusions, though their potential for contributing growth-promoting force remains unexplored. Using a novel inducible system that docks myosin motor domains to membrane binding modules with temporal control, we found that the application of myosin-generated force to the plasma membrane is sufficient for driving robust elongation of protrusions. Protrusion growth scaled with motor accumulation, required active, barbed end-directed force, and was independent of cargo delivery or the recruitment of canonical barbed end elongation factors. Application of growth-promoting force was also supported by structurally distinct myosin motor domains and membrane binding modules. We conclude that myosin-generated force can drive protrusion growth and this mechanism is likely active in diverse biological contexts.

## INTRODUCTION

Actin-based membrane protrusions are ancient cell surface features that have evolved to perform a vast array of biological roles needed for cell function and survival. Microvilli, stereocilia, and filopodia represent a group of structurally related protrusions supported by a core bundle of actin-filaments (Mooseker and Tilney, 1975; Small, 1988; Tilney et al., 1980). Despite a common morphology, these structures serve distinct physiological functions: microvilli drive nutrient absorption in the gut, stereocilia regulate hearing and balance in the ear, and filopodia contribute to a range of functions including cell migration, cell-cell adhesion, and neurite outgrowth (Houdusse and Titus, 2021). While the function, structure and composition of actin-based protrusions have been studied extensively for decades, the physical mechanisms that control their assembly are still emerging.

A key step in protrusion growth is deformation of the plasma membrane. Previous theoretical and experimental work suggested that positive (i.e. outward) membrane curvature is driven by the mechanical force generated during actin monomer incorporation at the membrane-proximal barbed ends of growing filaments (Footer et al., 2007; Hill and Kirschner, 1982a, b; Kovar and Pollard, 2004; Mogilner and Oster, 1996; Mogilner and Rubinstein, 2005; Peskin et al., 1993; Theriot, 2000). Elongation factors such as formin family proteins and enabled/vasodilator-stimulated phosphoprotein (Ena/VASP) have also been implicated in accelerating protrusion elongation, potentially by promoting rapid and processive incorporation of monomers at the membrane interface (Bear et al., 2002; Higashida et al., 2004; Kovar and Pollard, 2004). However, a recent study found that protrusion length is not tightly coupled to the levels of these canonical elongation factors (Dobramysl et al., 2021), highlighting the ambiguity of mechanisms that drive growth.

Although actin filament polymerization can drive protrusion growth, the potential for other force generators to contribute to this process has remained unclear. Indeed, myosin superfamily members are some of the most abundant residents of actin-based protrusions and may hold the potential to generate significant growth-promoting force (Jacquemet et al., 2019; Krey et al., 2017; McConnell et al., 2011). All myosins consist of three structural domains: a conserved N-terminal motor domain that binds actin and hydrolyzes ATP, a central neck that serves as a lever arm to amplify motion produced during the power stroke (Rayment et al., 1993; Uyeda et al., 1996; Warshaw et al., 2000), and a class-specific C-terminal tail domain (Sellers, 2000). Myosin tail domains mediate cargo binding and association with the plasma membrane (either directly or indirectly). Of all myosin classes, protrusions contain primarily barbed end directed-myosins: microvilli contain myosin-1a, -1d, and -7b (Benesh et al., 2010; Chen et al., 2001; Conzelman and Mooseker, 1987), stereocilia contain myosin-7a, -3a, and -15a (Belyantseva et al., 2003; Hasson et al., 1995; Schneider et al., 2006), and filopodia contain myosin-7 (Petersen et al., 2016) and myosin-10 (Berg et al., 2000). Several of these motors accumulate in the distal tip compartment, which presumably reflects their robust barbed end-directed movement in these systems. Consistent with this view, previous studies implicate myosins in the delivery of critical regulatory and structural cargoes to the distal tips, suggesting that these motors are important regulators of growth (Houdusse and Titus, 2021). Indeed, loss-of-function studies with tip-targeting myosins generally result in a loss of distal tip components and shorter protrusions in all three systems (Belyantseva et al., 2003; Bohil et al., 2006; Weck et al., 2016).

Beyond interacting directly with cargo proteins, the tail domains of myosins also interact with the plasma membrane, either directly or indirectly. Some myosins bind directly to the inner leaflet of the membrane through non-specific electrostatic interaction mediated by their highly basic C-terminal tail (Feeser et al., 2010; Mazerik and Tyska, 2012). Others interact with the plasma membrane using motifs that bind to specific membrane lipid head groups (Hokanson and Ostap, 2006; Plantard et al., 2010), or through interactions with membrane-associated peripheral or transmembrane proteins (Crawley et al., 2014; Yu et al., 2017; Zhang et al., 2004). From this perspective, myosin motors are well-positioned to exert significant force on the membrane, and consequently, impact the kinetics of protrusion growth. However, this hypothesis remains untested due to the technical challenge of uncoupling the contributions of tipward cargo transport from myosin tail membrane interactions.

To address this challenge and determine if membrane-bound myosin-generated force promotes protrusion growth, we developed a synthetic cell-based system that allows for temporally controlled docking of myosin motor domains (lacking cargo binding motifs) to the membrane. Our studies initially leverage the motor domain from the well-studied filopodia resident motor, myosin-10 (Myo10). Myo10 walks towards the barbed ends of polarized actin filament bundles, enriches at the distal tips of filopodia, and is sufficient for the initiation of filopodia formation and subsequent elongation (Berg and Cheney, 2002). Moreover, the Myo10 tail binds to cargoes (Tokuo and Ikebe, 2004) that are critical for filopodial function, but also interacts with the plasma membrane using a several distinct mechanisms (Plantard et al., 2010; Zhang et al., 2004). We found that membrane-bound Myo10 motor domains drive robust cell surface protrusion elongation independently of cargo binding and delivery activities. Live-cell and super-resolution imaging studies indicate that while the induced protrusions share many common features with canonical filopodia, they elongate independent of the formin, mammalian diaphanous-related formin 1 (mDia1), and VASP, suggesting myosin-generated force circumnavigates the usual requirement for these factors. Motor domain-driven filopodial elongation was also supported by a variety of membrane binding motifs, although integral membrane motifs produced the most robust response. Elongation activity was not specifically dependent on the mechanical properties of Myo10, as the motor domains from stereocilia motors, myosin-15a and -3a, also promoted filopodia growth. To our knowledge, these data are the first to reveal that application of myosin-generated force to the plasma membrane directly drives protrusion growth.

## RESULTS

### Membrane-bound Myo10 motor domains drive robust protrusion elongation, independent of cargo binding

To investigate the role of myosin-generated force in protrusion growth, we used a rapalog inducible system that offers switchable control over the oligomerization of proteins tagged with FRB and FKBP (Banaszynski et al., 2005; Inobe and Nukina, 2016). These modules were used to engineer constructs that enabled us to rapidly induce interaction between myosin motor domains (tagged with FRB) and plasma membrane binding motifs (tagged with FKBP) upon the addition of rapalog (**Fig 1A**). We first examined the effect of docking the motor domain from the filopodial motor, Myo10, to the plasma membrane in live HeLa cells. To eliminate cargo binding and specifically probe how the application of force to the membrane affects protrusion growth, we replaced the entire C-terminal tail of Myo10 with

**Figure 1:**
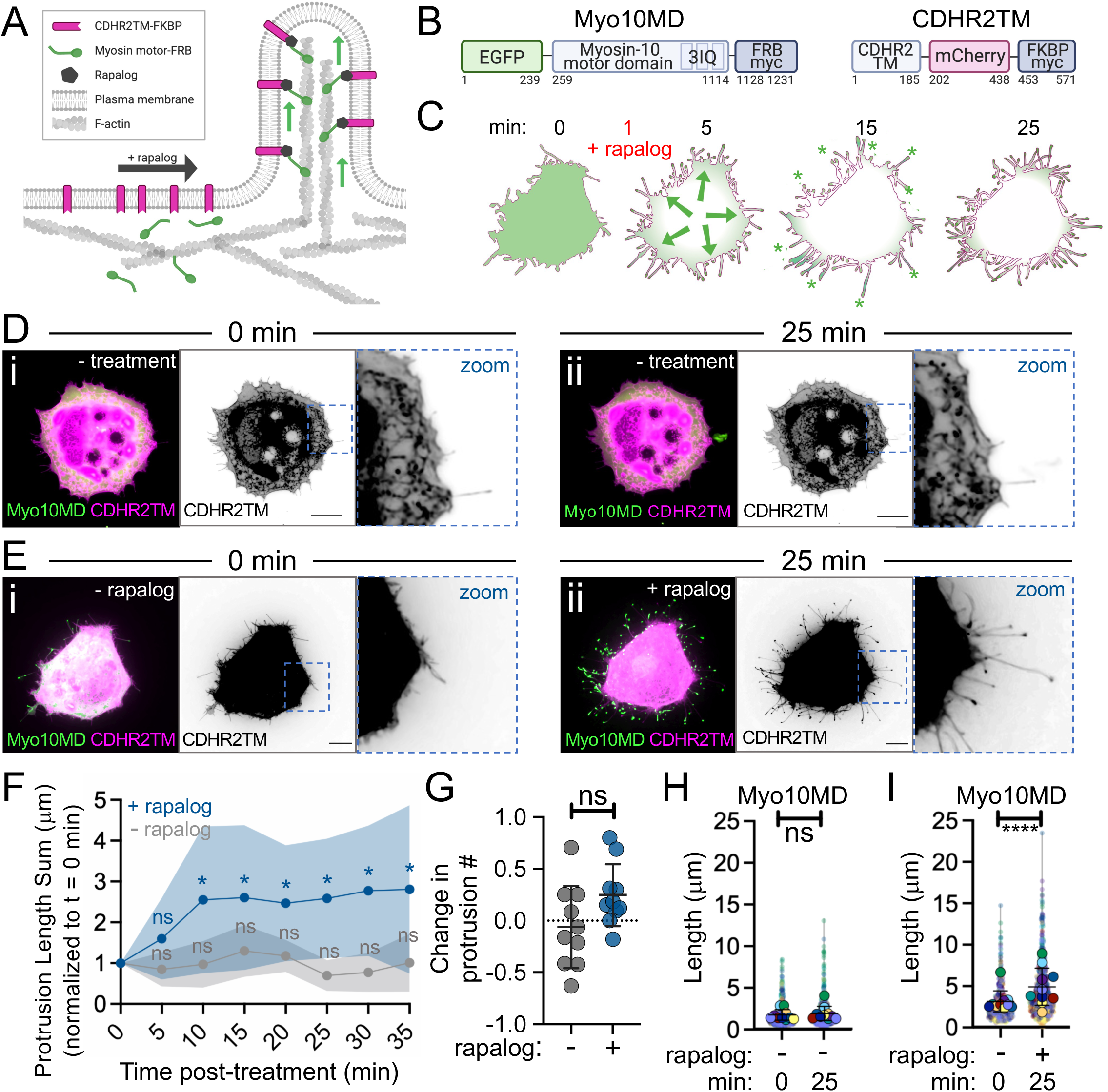
Membrane bound myosin-10 motor domains drive protrusion elongation. (**A**) Cartoon of the rapalog inducible system used to dock the myosin motor domains onto the plasma membrane. (**B**) Cartoons depicting the CDHR2TM and Myo10MD constructs for these experiments; numbers represent amino acids. (**C**) Cartoon depicting the timeline of rapalog addition and the resultant relocalization of the Myo10MD (green) construct to the CDHR2TM-labeled plasma membrane (magenta). Green asterisks mark Myo10MD enrichment at the tips of protrusions. (**D**) Confocal maximum intensity projection images of control HeLa cells expressing Myo10MD (green) and CDHR2TM (magenta) at 0 min (**i**) and 25 min (**ii**) with no rapalog treatment. (**E**) Confocal maximum intensity projection of HeLa cells expressing Myo10MD (green) and CDHR2TM (magenta) at 0 min (**i**) and 25 min after (**ii**) rapalog treatment. (**F**) Length sum of protrusions over time in untreated (gray) or rapalog treated (blue) HeLa cells expressing Myo10MD and CDHR2TM; *n* = 8 individual cells for each treatment. (**G**) Fractional change in protrusion number from 0 min to 25 min in untreated (gray) or rapalog treated (blue) HeLa cells expressing Myo10MD and CDHR2TM; *n* = 10 individual cells for each condition. (**H**) Length of individual protrusions measured at 0 min and 25 min in untreated HeLa cells expressing Myo10MD and CDHR2TM; *n* = 10 individual cells, *n* = > 650 individual protrusions. (**I**) Length of individual protrusions measured at 0 min and 25 min post rapalog treatment in HeLa cells expressing Myo10MD and CDHR2TM*; n* = 10 individual cells, *n* = > 550 individual protrusions. Data in **(H**) and (**I**) are represented as a Superplot, where transparent circles represent the length of individual protrusions, opaque circles represent the average length of protrusions of individual cells, violin plots show the distribution of the data, and all color matched circles represent measurements from the same cell. All graph error bars represent the mean ± SD; ns, p-value > 0.05, * p-value ≤ 0.05, **** p-value ≤ 0.0001. Scale bars = 10 μm.

FRB (a.a. 1-855, EGFP-Myo10MD-FRB, herein referred to as Myo10MD) (**Fig 1B**). For a membrane-binding motif, we used the transmembrane domain (TM; a.a. 1155-1310) from cadherin related family member 2 (CDHR2), a single-spanning membrane protein not endogenously found in HeLa cells (CDHR2TM-mCherry-FKBP, herein referred to as CDHR2TM) (**Fig 1B**). Using live-cell spinning-disc confocal (SDC) microscopy, we visualized cells expressing both Myo10MD and CDHR2TM, imaging every 5 minutes before and after the addition of rapalog (**Fig 1C**). Upon rapalog treatment, we noticed clear relocalization of the Myo10MD signal from the cytoplasm to the cell periphery, as expected based on the plasma membrane localization of CDHR2TM. Strikingly, cells displayed a robust elongation of finger-like protrusions within 10 minutes of rapalog treatment compared to untreated control cells (**Fig 1D, 1E - compare Fig 1Dii to 1Eii- Fig 1F, Video S1, S2**). While we were able to directly visualize *de novo* protrusion growth events in our time-lapse data, and the number of protrusions did trend upward, this increase was not significant (**Fig 1G**). However, protrusion length increased significantly when Myo10MD was docked to the membrane (**Fig 1H, 1I**), suggesting that myosin-generated forces contribute primarily to elongating existing structures rather than initiating new ones.

To determine if protrusion growth induced by membrane-bound Myo10MD was unique to HeLa cells, we co-expressed our system components in B16F1 *Mus musculus* melanoma cells, which also form fascin-bundled filopodia, and LLC-PK1-CL4 (CL4) *Sus scrofa* kidney cells, which form microvilli on their apical surface. Induction of Myo10MD membrane binding in both cell lines drove robust protrusion elongation. Thus, the application of myosin-generated force to the plasma membrane is sufficient for driving protrusion growth in distinct cell types derived from a range of tissues (**Fig. S1**).

### Protrusions induced by membrane-bound Myo10MD exhibit features of filopodia

We next sought to determine if the protrusions induced by membrane-bound Myo10MD represent canonical filopodia. To this end, HeLa cells were fixed 25 minutes after the addition of rapalog, and then stained for F-actin (phalloidin)(Vandekerckhove et al., 1985) and the filopodia-specific parallel filament bundler, fascin (Vignjevic et al., 2006). Super-resolution structured illumination microscopy (SIM) of these samples revealed that protrusions induced by membrane-bound Myo10MD contained both F-actin and fascin (**Fig. 2A, B, S2)**. We also analyzed SDC time-lapse data and determined that individual protrusions induced by membrane bound Myo10MD elongated at a similar rate to canonical filopodia. Individual protrusions induced by membrane-bound Myo10MD elongated at 2.16 μm/min, a rate similar to that reported for filopodia in previous studies (∼2.2 μm/min)(Schäfer et al., 2011)(**Fig 2C, 2Ci zoom, 2D**). Based on current models of filopodial growth, we also expected that protrusion growth induced by membrane-bound Myo10MD would require actin polymerization. Indeed, HeLa cells treated with the actin poisons cytochalasin D (caps filament barbed ends) and latrunculin A (sequesters actin monomers), exhibited significantly reduced protrusion elongation in response to rapalog induction (**Fig S3A, B**). Thus, protrusions induced by membrane-bound Myo10MD are similar to canonical filopodia in that they are supported by parallel, fascin-bundled actin filaments, and require both uncapped barbed ends and a pool of free actin monomers to elongate.

**Figure 2:**
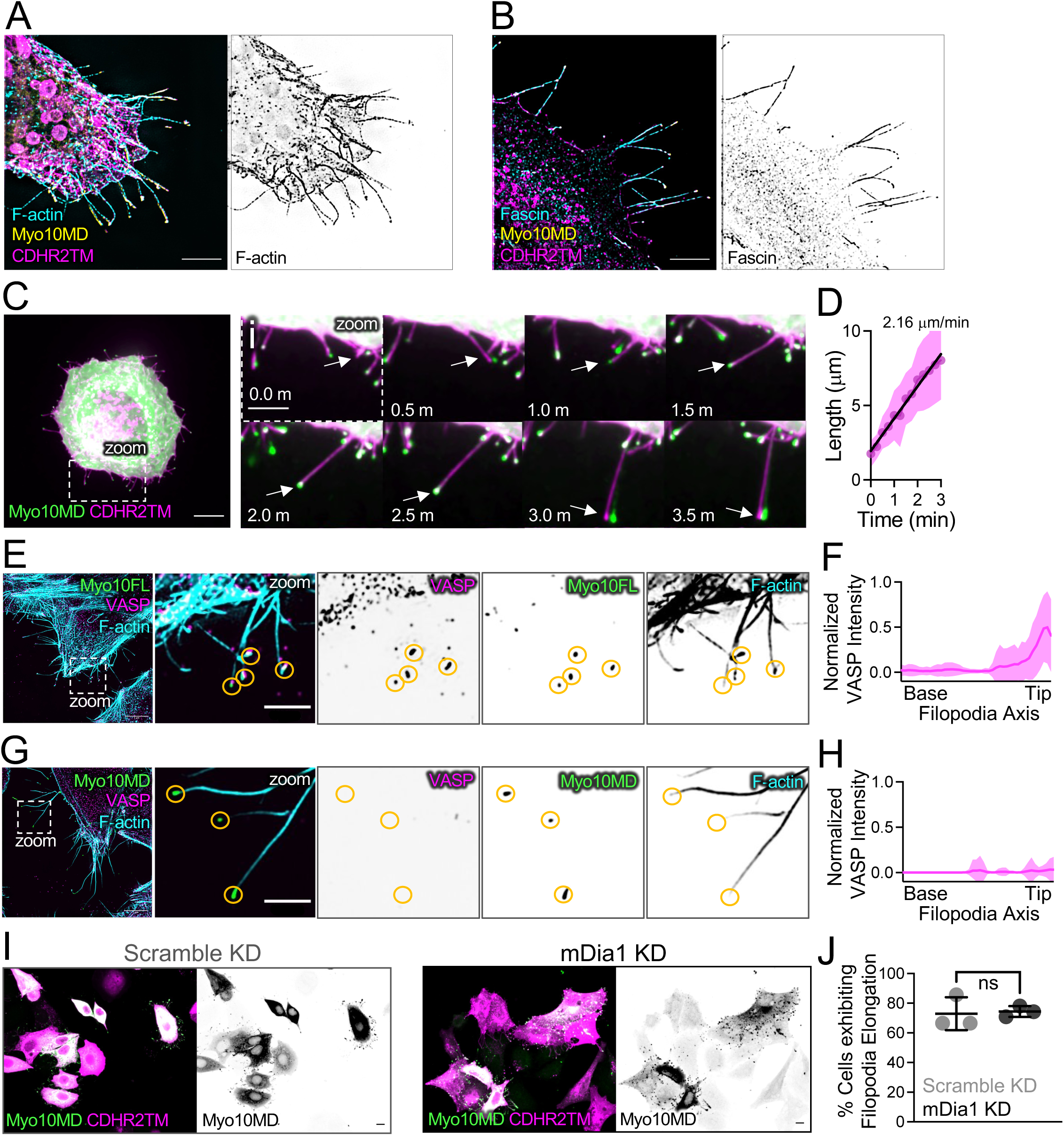
Protrusions induced by membrane bound Myo10MD exhibit features of filopodia. (**A**) SIM maximum intensity projection image of a HeLa cell expressing Myo10MD (yellow) and CDHR2TM (magenta) and stained for F-actin with phalloidin (cyan). Panel to the right shows inverted single channel image of F-actin (phalloidin). Scale bar = 5 μm. (**B**) SIM maximum intensity projection image of HeLa cells expressing Myo10MD (yellow) and CDHR2TM (magenta) and stained for endogenous fascin (cyan). Panel to the right shows inverted single channel image of fascin. Scale bar = 5 μm. (**C**) Confocal maximum intensity projection of HeLa cell expressing Myo10MD (green) and CDHR2TM (magenta) 2 min after rapalog addition. Scale bar = 10 μm. (**Ci**) Montage of zoomed in region (**C**) highlighting the growth of a single filopodium (white arrow). (**D**) Elongation rate of single filopodium induced by rapalog treatment. Rate was calculated via the slope using a simple linear regression; *n* = 14 individual filopodia from *n* = 3 separate cells. (**E**) SIM maximum intensity projection image of a HeLa cell expressing full-length Myosin-10 (green) and stained for endogenous VASP (magenta) and F-actin with phalloidin (cyan). Zoom images to the right show merge and inverted single channel images. Yellow circles denote the tips of individual filopodia. (**F**) Line scans parallel to the filopodial axis show the intensity distribution of VASP in filopodia generated from over-expressing EGFP-myosin-10 full length; *n* = 30 protrusions, line represents the average intensity. (**G**) SIM maximum intensity projection image of a HeLa cell expressing Myo10MD (green) and CDHR2TM (not shown) 25 min after rapalog treatment and stained for endogenous VASP (magenta) and F-actin (cyan). Zoom images to the right show merge and inverted single channel images. Yellow circles denote the tips of individual filopodia. For **E** and **G**, scale bar = 5 μm, zoom scale bar = 2 μm. (**H**) Line scans parallel to the filopodial axis show the intensity distribution of VASP in filopodia; *n* = 39 filopodia, line represents the average intensity. (**I**) Confocal maximum intensity projection of HeLa cells transfected with either non-targeting siRNA (scramble KD) or siRNA targeting mDia (mDia1 KD) and expressing Myo10MD (green) and CDHR2TM (magenta), imaged 25 min after rapalog treatment. Scale bars = 10 μm. (**J**) Percent of cells displaying filopodial elongation with or without KD of mDia1; *n* = 3 replicates with 87-93% knockdown efficiency. Double transfected cells/condition scramble KD (*n* = 184) and mDia1 KD (*n* = 63). Ns, p-value > 0.05 calculated using the unpaired Student’s t-test; all graph error bars represent the mean ± SD.

### Membrane-bound Myo10MD promotes filopodial growth independent of barbed end elongation factors

Previous studies identified barbed end elongation factors as proteins that enrich at the distal ends of filopodia and in turn promote their growth. Ena/VASP and formin family proteins are well-studied examples (Bear et al., 2002; Higashida et al., 2004) and we sought to determine if membrane-bound Myo10MD promoted filopodial growth by recruiting these factors. We first examined HeLa cells for the enrichment of Ena/VASP at the tips of induced filopodia. Ena/VASP is a tetrameric protein that promotes actin monomer incorporation into filaments and protects barbed ends from capping protein, which would otherwise stall core bundle elongation. VASP binds to full-length Myo10 and may be transported to the distal ends of filopodia via this interaction (Tokuo and Ikebe, 2004), although the possibility of such direct transport is eliminated in our experimental system as Myo10MD constructs lack the C-terminal tail. Using SIM, we visualized cells transfected with either full-length Myo10 or the combination of Myo10MD and CDHR2TM that drives filopodial induction. As expected, filopodia generated by over-expressing full length Myo10 displayed prominent VASP enrichment at their distal tips (**Fig 2E, 2F**).

Remarkably, VASP was undetectable at the tips of filopodia induced by membrane-bound Myo10MD (**Fig 2G, 2H**). We also asked if formin family proteins were involved in the rapalog-induced response. Based on RNAseq data (Uhlen et al., 2015), mDia1 is the most highly expressed formin in HeLa cells. Moreover, previous studies implicate mDia1 as a processive elongator of actin filaments in filopodial core bundles (Higashida et al., 2004). To examine potential involvement of mDia1, we knocked down its expression in HeLa cells using siRNA (**Fig S4**), transfected cells with Myo10MD and CDHR2TM, and induced filopodial growth with rapalog treatment. Interestingly, we observed no change in the fraction of cells that were able to elongate filopodia in mDia1 KD cells relative to controls (**Fig 2I, 2J**). Together, these results suggest that docking Myo10MD force on the plasma membrane is sufficient to drive filopodial elongation, independent of the canonical barbed end elongation factors VASP and mDia1.

### Filopodial elongation temporally correlates with Myo10MD accumulation at the distal tips

We next sought to determine how membrane-bound Myo10MD elongates filopodia in the absence of barbed end elongation factors. One possibility is that mechanical force applied by Myo10MD directly to the plasma membrane promotes growth by reducing the physical barrier to protrusion elongation. A testable prediction that emerges from this hypothesis is that active, barbed end-directed force generation by membrane-bound Myo10MD is required to induce filopodial growth in response to rapalog treatment. To test this, we first sought to determine if higher levels of Myo10MD, and therefore, higher levels of force, might drive faster elongation rates and/or longer protrusions. While the absolute amount of force generated by Myo10MD in cells is experimentally inaccessible, we reasoned that we could obtain a relative measure of force by monitoring the EGFP intensity from the Myo10MD construct; this signal is proportional to the number of Myo10MD motors and thus local force-generating potential. Using live-cell SDC imaging, we monitored Myo10MD tip intensity during individual filopodial growth events after inducing HeLa cells with rapalog. Because filopodia generated in these experiments are highly dynamic and poorly anchored to the coverslip substrate, culture media was supplemented with 0.5% methylcellulose; this prevented lateral waving of protrusions and improved our ability to monitor the details of elongation with high temporal resolution. Importantly, the addition of methylcellulose did not alter the growth rates of filopodia induced by membrane-bound Myo10MD (**Fig S5**). Using this approach, we measured the distal tip intensity of Myo10MD in parallel with filopodial length from time-lapse data collected at 2 sec intervals (**Fig 3A, Video S3**). This analysis revealed a striking and direct temporal correlation between the length of a growing filopodium and the accumulation of Myo10MD at its distal tip (**Fig 3B**). Thus, during growth, filopodial length is well correlated to the force-generating potential of tip-enriched myosin motors.

**Figure 3:**
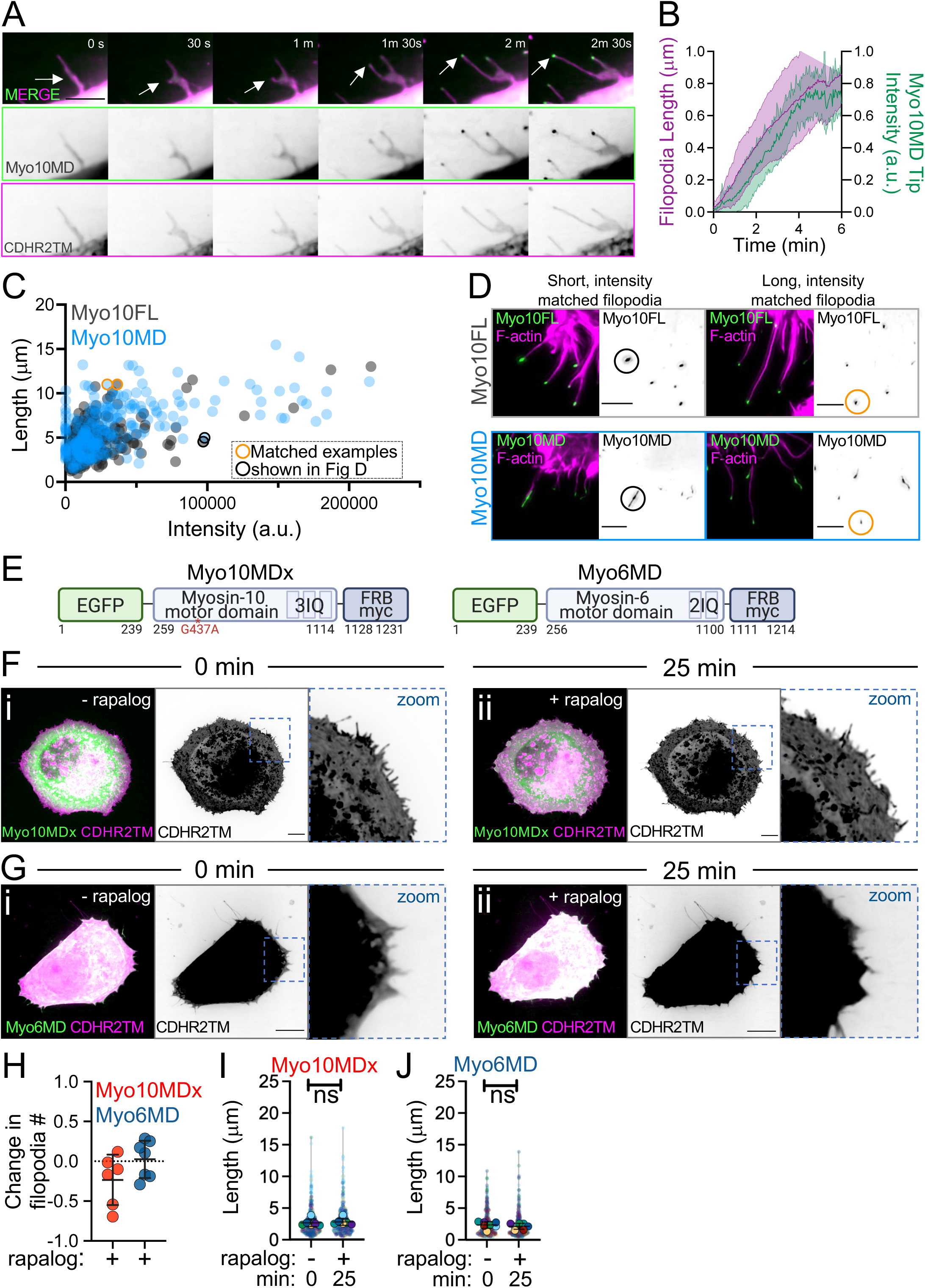
Barbed end-directed force is required for filopodial induction by Myo10MD. (**A**) Montage of a filopodial growth event from a HeLa cell expressing Myo10MD (green) and CDHR2TM (magenta) imaged in 0.5% methylcellulose. Arrows denote the tip of a single, growing filopodium. Scale bar = 5 μm. (**B**) Normalized length (magenta) and tip intensity (green) vs. time curves for individual filopodia on cells expressing Myo10MD and CDHR2TM; *n* = 10 filopodia. (**C**) Filopodial length and myosin distal tip-intensity from individual protrusions in cells expressing Myo10FL (gray) or Myo10MD (blue); each circle represents an individual filopodium. Orange and blue circled data points represent filopodia shown in **D**; *n* = 304 filopodia for Myo10FL and 273 for Myo10MD. (**D**) Confocal maximum intensity projections of length and tip intensity-matched filopodia in cells expressing Myo10FL or Myo10MD (green), and stained for F-actin with phalloidin (magenta); single inverted channel images are shown to the right. Scale bar = 5 μm. (**E**) Cartoons depicting the myosin-10 motor domain dead G437A mutant (Myo10MDx) and pointed end-directed myosin-6 motor domain (Myo6MD) constructs; numbers below represent amino acids. (**F**) Confocal maximum intensity projection of HeLa cells expressing Myo10MDx (green) and CDHR2TM (magenta) before and 25 min after (**ii**) rapalog treatment. (**G**) Confocal maximum intensity projection of HeLa cells expressing Myo6MD (green) and CDHR2TM (magenta) before (**i**) and 25 min after (**ii**) rapalog treatment. For **F** and **G**, scale bar = 10 μm. (**H**) Fractional change in filopodia number in HeLa cells expressing CDHR2TM and Myo10MDx (red) or Myo6MD (blue) 0 min and 25 min after rapalog treatment; *n* = 6-7 individual cells for each condition. (**iii**) Filopodial length measured at 0 min and 25 min post rapalog treatment in HeLa cells expressing Myo10MDx and CDHR2TM; *n* = 6 individual cells, *n* = > 420 individual protrusions. (**J**) Filopodial length measured at 0 min and 25 min post rapalog treatment in HeLa cells expressing Myo6MD and CDHR2TM; *n* = 7 individual cells, *n* = > 365 individual protrusions. Data in (**I**) and (**J**) are represented as a Superplot, where transparent circles represent the length of individual filopodia, opaque circles represent the average length of filopodia from individual cells, violin plots show the distribution of the data, and all color-matched circles represent measurements from the same cell. All graph error bars represent the mean ± SD; ns, p-value > 0.05.

To place these findings into context and determine how the distal tip enrichment of Myo10MD compares to wild type full-length Myo10 (Myo10FL), we repeated these experiments in HeLa cells expressing EGFP-Myo10FL and scored tip intensities and lengths from filopodia. This analysis revealed that the tip intensity vs. length distributions obtained for Myo10MD and Myo10FL largely overlapped (**Fig 3C**). Moreover, filopodia generated by both constructs had similar morphology when matched in length and intensity (**Fig 3D - circled data points in 3C**). Together these data indicate that, during filopodial elongation, the tip enrichment and thus force-generating potential of membrane-bound Myo10MD is comparable to Myo10FL.

### Active, barbed end-directed force is required for filopodial induction

The observation that filopodial length increases in parallel with distal tip accumulation of membrane-bound Myo10MD is consistent with the proposal that force generated by this motor directly promotes protrusion growth. From this perspective, we next examined whether the mechanical activity of Myo10MD is required for filopodia elongation. We generated two additional Myo10MD constructs containing mutations in switch I (R220A) or switch II (G437A) that are expected to disrupt force generation (Sasaki and Sutoh, 1998). Docking the motor-dead G437A construct (EGFP-Myo10MDx-FRB, herein referred to as Myo10MDx) (**Fig 3E**) to the plasma membrane with rapalog treatment failed to elongate filopodia (**Fig 3Fi-ii, 3H, 3I, Video S4**). A motor-dead R220A Myo10MD mutant also failed to accumulate on the plasma membrane or in filopodia, most likely because of its impaired actin binding (**Fig S6**). Based on these data, we conclude that active, force-generating motors are needed to drive filopodial elongation in response to rapalog treatment.

To determine if the induction of filopodial elongation specifically requires the application of barbed end-directed force to the plasma membrane, we generated a construct using the motor domain from myosin-6, the sole pointed end-directed member of the myosin superfamily (a.a. 1-844, EGFP-Myo6MD-FRB, herein referred to as Myo6MD) (**Fig 3E**). Membrane-bound Myo6MD also failed to promote filopodial elongation in response to rapalog treatment (**3Gi-ii, 3H, 3J, Video S5**). Taken together, these data indicate that the application of barbed end-directed force by membrane-bound motors is needed for robust filopodia elongation.

### Filopodial growth induced by membrane-bound Myo10MD is limited by the availability of actin monomers and plasma membrane

Why the robust filopodial growth induced by membrane-bound Myo10MD stalls at long lengths remains unclear. One possibility is that protrusions become so long that the pool of free actin monomers available for incorporation at the distal barbed ends falls below the level needed to continue elongation. To test this possibility, HeLa cells were subjected to long-term G-actin sequestration with a low concentration of Latrunculin A (Lat A). Our rationale was that washing out Lat A post-rapalog treatment would rapidly and acutely increase the pool of free actin monomers; any “reactivation” of filopodial growth at that point would indicate that the availability of free G-actin normally limits elongation. For these experiments, HeLa cells were incubated in 100 nM Lat A for 7 days, to allow for long-term adaption prior to transfection with Myo10MD and CDHR2TM (**Fig 4A**). Remarkably, Lat A washout induced a modest but detectable elongation of filopodia beyond their initial length (35 min) following rapalog treatment (**Fig 4B, C, compare blue and gray lines at 75 min, Video S6, S7**). These results suggest that the actin monomer pool does become limiting, and that the sustained elongation of protrusions induced by membrane-bound Myo10MD may be at least partially dependent on available monomers. Filopodial growth after rapalog induction might also be limited by the availability of encapsulating plasma membrane lipids (Raucher and Sheetz, 2000). To test this idea, we acutely expanded plasma membrane surface area by introducing CellMask-Orange (herein referred to as CellMask) to cells 35 min after filopodial elongation was initiated with rapalog treatment (**Fig 4D**). Interestingly, the addition of CellMask induced a second stage of elongation beyond the initial elongation driven by membrane-bound Myo10MD (**Fig 4E, F, compare magenta and gray lines at 140 min, Video S8, S9**). Thus, in addition to being limited by the availability of actin monomers, sustained filopodial growth driven by membrane-bound Myo10MD may also be limited by the availability of encapsulating plasma membrane.

**Figure 4:**
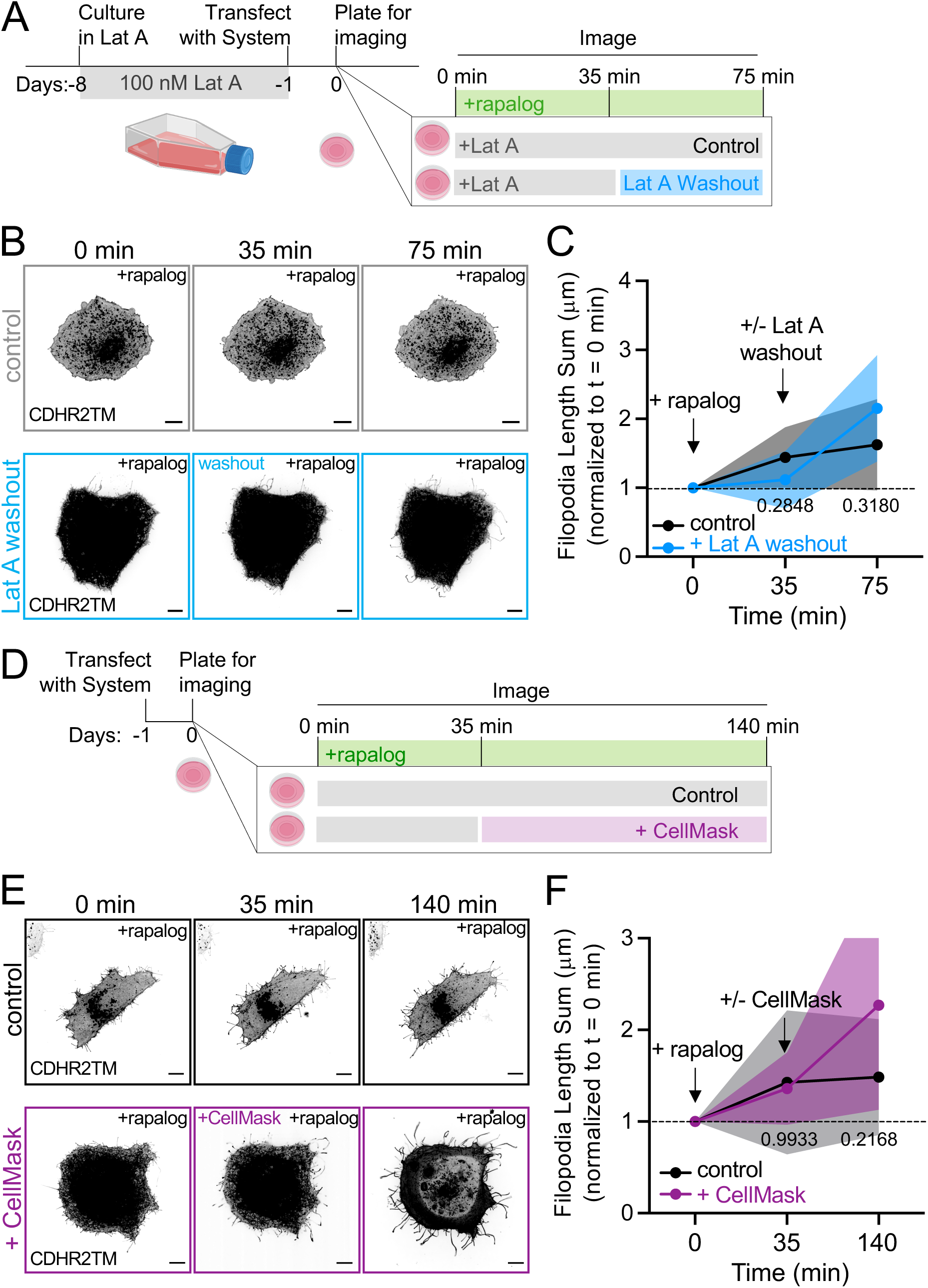
Myo10MD induced filopodial growth depends on membrane and G-actin availability. (**A**) Cartoon of the Latrunculin A (Lat A) washout experimental timeline. (**B**) Confocal maximum intensity projection of representative HeLa cells expressing Myo10MD (not shown) and CDHR2TM (gray scale), treated with 100 nM Lat A for one week (top; control) or Lat A for one week followed by washout (bottom). (**C**) Normalized length sum of filopodia generated over time in cells treated with Lat A (gray) or treated with Lat A followed by washout (blue). (**D**) Cartoon of membrane expansion experimental timeline. (**E**) Confocal maximum intensity projection of representative HeLa cells expressing Myo10MD (not shown) and CDHR2TM (gray scale). Untreated control cell (top) and cell treated with membrane intercalating CellMask for 35 min (bottom). (**F**) Normalized length sum of filopodia generated over time in control HeLa cells (gray) or cells treated with CellMask (magenta). All graph error bars are mean ± SD. P-values were calculated using a two-way ANOVA comparing treatments at the indicated time point, *n* = 10 individual cells for each of the four treatments. Scale bars = 10 μm.

### Myosin motor domains from structurally diverse superfamily classes also support rapalog-induced filopodial growth

Myosin superfamily members exhibit a range of mechanochemical properties, which enable these motors to perform specific subcellular tasks that hold unique kinetic requirements (e.g., cargo anchoring vs. transport). Therefore, we next sought to determine if the robust filopodial elongation driven by Myo10MD is unique to this motor, or could instead be supported by other myosins typically found in other actin-based protrusions, such as microvilli and stereocilia. To test this, we generated additional constructs based on protrusion-resident barbed end-directed myosins that exhibit a range of mechanochemical properties including myosin-1a, myosin-5b (lever arm with 6 IQ motifs (-wildtype) or 3 IQ motifs (-5b3IQ)), myosin-7b, myosin-3a, myosin-15a. By surveying fixed and stained samples after induction, we noted that the motor domains of myosin-1a, -5b-wildtype, -5b3IQ, and -7b were unable to significantly elongate filopodia when docked onto the plasma membrane (**Fig S7**). In contrast, motor domains from the stereocilia tip-targeting motors, myosin-3a (a.a. 407-1432, EGFP-Myo3aΔKMD-FRB, herein referred to as Myo3aΔKMD; **Fig 5A**) and myosin-15a (a.a. 1-750, EGFP-Myo15aMD-FRB, herein referred to as Myo15aMD; **Fig 5F**) drove significant elongation following rapalog induction (**Fig 5Bi, Bii, Gi, Gii, Video S10, S11)**. Quantification of the length sum of filopodia over time showed that there was a significant increase generated by both motors after 20 and 10 minutes of rapalog treatment, respectively (**Fig 5C, H**). Like the Myo10MD construct, docking either the Myo3aΔKMD or Myo15aMD constructs to the membrane did not significantly change the number of filopodia generated post-rapalog treatment (**Fig 5D, I**), but did increase filopodial length (**Fig 5E, J**). These results indicate that the robust filopodial elongation observed following rapalog induction is not unique to Myo10MD and can also driven by other structurally diverse myosins that reside in distinct protrusion environments.

**Figure 5:**
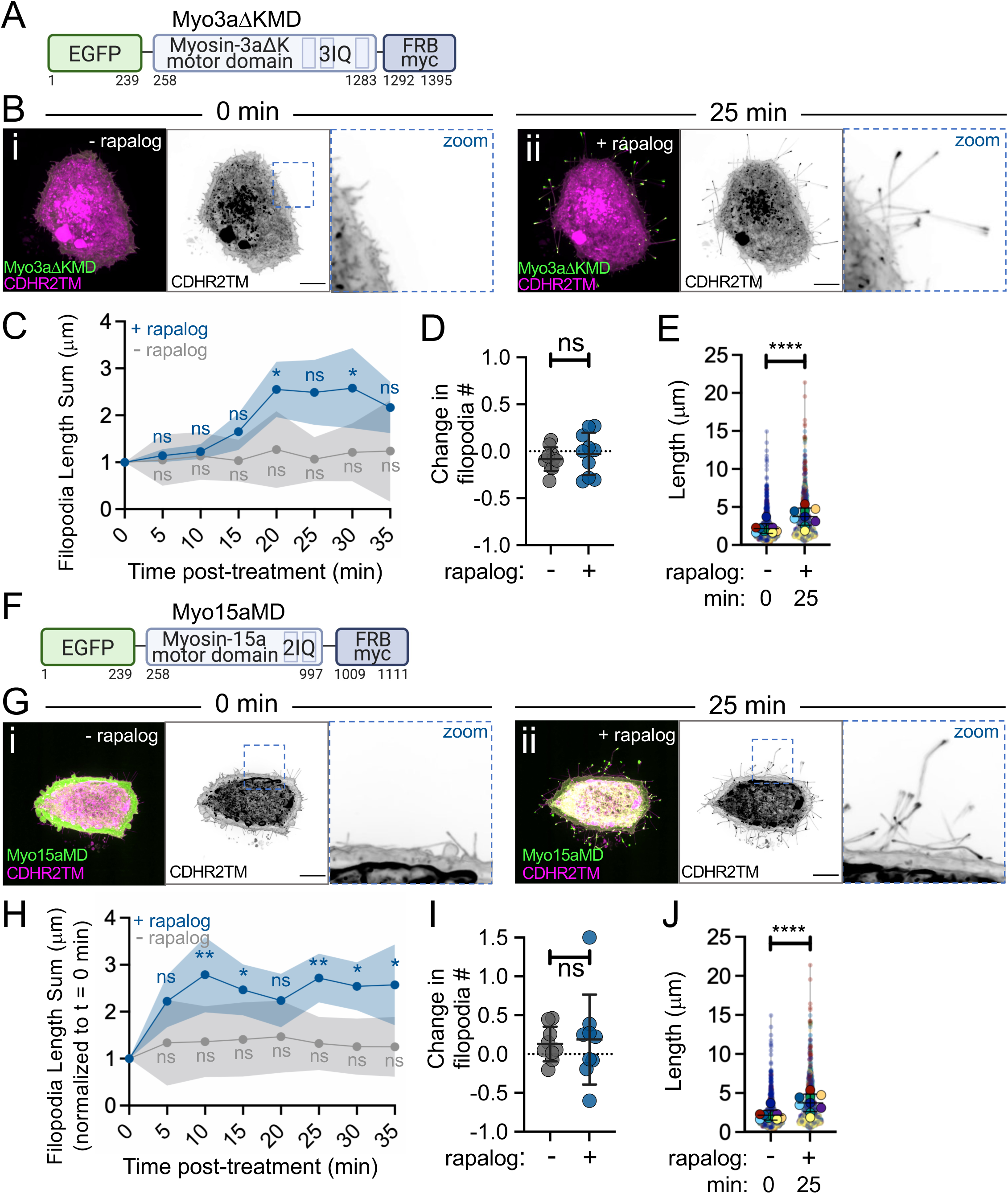
Motor domains from Myo3a and Myo15a also promote filopodia growth. (**A**) Cartoon depicting the EGFP-Myo3aΔK motor domain-FRB-myc construct (Myo3aΔKMD); numbers below represent amino acids. (**B**) Confocal maximum intensity projection image of control HeLa cells expressing the Myo3aΔKMD (green) and CDHR2TM (magenta) at 0 min (**i**) and 25 min after (**ii**) rapalog treatment. (**C**) Length sum of filopodia over time in untreated (gray line) or rapalog-treated (blue line) HeLa cells expressing Myo3aΔKMD and CDHR2TM; *n* = 10 cells per treatment. (**D**) Fractional change in filopodia number from 0 min to 25 min in untreated (gray) or rapalog treated (blue) HeLa cells expressing Myo3aΔKMD and CDHR2TM; *n* = 10 cells per condition. (**E**) Filopodia length at 0 and 25 min post rapalog treatment in HeLa cells expressing Myo3aΔKMD and CDHR2TM; *n* = 9 cells, *n* = > 670 protrusions. (**F**) Cartoon depicting the EGFP-Myosin-15a motor domain-FRB-myc construct (Myo15aMD); numbers below represent amino acids. (**G**) Confocal maximum intensity projection images of control HeLa cells expressing Myo15aMD (green) and CDHR2TM (magenta) before (**i**) and 25 min after (**ii**) rapalog treatment. (**H**) Length sum of filopodia over time in untreated (gray line) or rapalog treated (blue line) HeLa cells expressing Myo15aMD and CDHR2TM; *n* = 10-11 individual cells per treatment. (**I**) Fractional change in filopodial number from 0 to 25 min in untreated (gray) or rapalog treated (blue) HeLa cells expressing Myo15aMD and CDHR2TM; *n* = 9 cells per condition. (**J**) Filopodia length measured at 0 and 25 min post rapalog treatment in HeLa cells expressing Myo15aMD and CDHR2TM; *n* = 9 individual cells, *n* = > 730 protrusions. Data in (**E**) and (**J**) are represented as a Superplot, where transparent circles represent the length of individual filopodia, opaque circles represent the average length of protrusions of individual cells, violin plots show the distribution of the data, and all color matched circles represent measurements from the same cell. All graph error bars represent the mean ± SD; ns p-value > 0.05, * p-value ≤ 0.05, ** p-value ≤ 0.01, **** p-value ≤ 0.0001. Scale bars = 10 μm.

### Myo10MD driven filopodial elongation is supported by diverse membrane binding motifs

Our data show that docking the Myo10MD on the plasma membrane using the single-spanning CDHR2TM leads to robust filopodial elongation. In addition to binding transmembrane proteins, some myosin tail domains interact directly with the plasma membrane. For example, the second PH domain in the myosin-10 tail interacts with the phosphoinositol species, PI(3,4,5)P_3_, which is found in the plasma membrane along the filopodia shaft, and the TH1 domain of myosin-1a interacts with the acidic phospholipids found in the inner leaflet of the plasma membrane. To determine if these peripheral modes of membrane binding support filopodial elongation, we generated two additional membrane docking constructs by fusing FKBP to: (1) the PH domain of Bruton’s tyrosine kinase (BTK), which binds PI(3,4,5)P_3_ (Fukuda et al., 1996; Salim et al., 1996)(BTK-PH-mCherry-FKBP, herein referred to as BTK-PH, **Fig 6D**), and (2) the TH1 domain of myosin-1a, which binds electrostatically to the inner leaflet of the plasma membrane (Mazerik and Tyska, 2012)(FKBP-mCherry-TH1, herein referred to as TH1; **Fig 6G**). Both constructs showed membrane enrichment as expected (**Fig 6B, E, H,** pre-bleach). Fluorescence recovery after photobleaching (FRAP) quantification of the turnover kinetics in each case also revealed distinct immobile fractions (I.F.) for each construct (**Fig 6C, F, I**). As predicted, the integral, transmembrane CDHR2TM construct (**Fig 6A**) had the largest I.F. compared to the two peripheral membrane-interacting constructs (compare 6C to 6F and 6I). To determine if these different membrane binding motifs could support filopodial elongation, we turned to a fixed-cell approach, where HeLa cells were transfected with either of the three membrane constructs and the Myo10MD construct and induced to produce filopodia with the addition of rapalog. This analysis showed that all three modes of membrane interactions were able to support filopodia elongation (**Fig 6J, K, L**), with the CDHR2TM producing the most robust response (**Fig 6M**). These results further suggest that the ability of myosin motor domains to drive filopodia elongation is independent of the structural details of membrane association.

**Figure 6:**
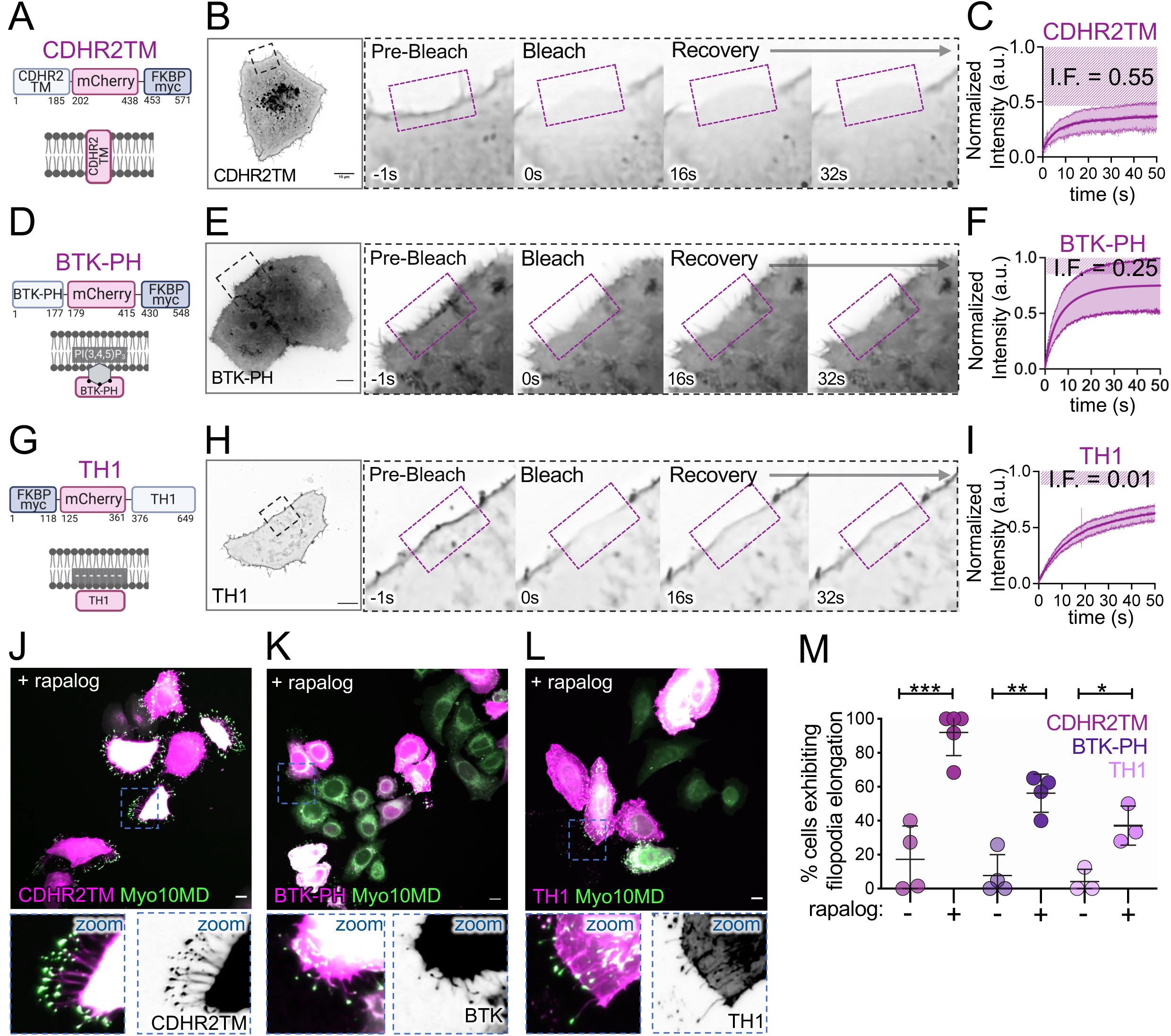
Myo10MD driven filopodial elongation is supported by diverse membrane binding motifs. (**A**) Cartoon depicting the integral transmembrane construct CDHR2TM-mCherry-FKBP construct. (**B**) Confocal maximum intensity projection timelapse montage showing fluorescence recovery after photobleaching (FRAP) analysis of a HeLa cell expressing CDHR2TM. (C) FRAP recovery curve; measurements were taken from ROI shown in B; *n* = 6 cells; magenta striped region indicates the immobile fraction. (**D**) Cartoon showing organization of the peripheral membrane binding construct BTK-PH- mCherry-FKBP (BTK-PH). (**E**) Confocal maximum intensity projection timelapse montage showing FRAP analysis of a HeLa cell expressing BTK-PH. (**F**) FRAP recovery curve; measurements were taken from the ROI shown in E; *n* = 9 cells; magenta striped region indicates the immobile fraction. (**G**) Cartoon depicting the peripheral membrane binding construct FKBP-mCherry-TH1 (TH1). (**H**) Confocal maximum intensity projection timelapse montage showing FRAP analysis of a HeLa cell expressing TH1. (I) FRAP recovery curve; measurements were taken from the ROI shown in H; *n* = 8 cells; magenta striped region indicates the immobile fraction. (**J**) Confocal maximum intensity projection image of HeLa cells expressing Myo10MD (green) and CDHR2TM (magenta) 25 min after rapalog treatment. (**K**) Confocal maximum intensity projection image of HeLa cells expressing Myo10MD (green) and BTK-PH (magenta) 25 min after rapalog treatment. (**L**) Confocal maximum intensity projection image of HeLa cells expressing Myo10MD (green) and TH1 (magenta) 25 min after rapalog treatment. (**M**) Percentage of untreated or rapalog-treated HeLa cells expressing Myo10MD and CDHR2TM, BTK-PH, or TH1 exhibiting filopodia elongation; *n* > 170 cells for CDHR2TM, *n* >140 cells for BTK-PH, and *n* > 220 cells for TH1. All graph error bars represent the mean ± SD; * p-value ≤ 0.05, ** p-value ≤ 0.01, *** p-value ≤ 0.001. Scale bars = 10 μm.

## DISCUSSION

Cell surface protrusions are essential for mediating physical and biochemical interactions with the external environment. Throughout evolution, μm-scale finger-like protrusions have been adapted to fill a range of functional niches, including mechanosensation and solute transport (Revenu et al., 2004). Fundamental to the formation of a surface protrusion is the outward deformation of the fluid plasma membrane. Estimates of the forces a cell must apply to form a protrusion come from optical trap-based measurements of the “tether force”: the force required to pull and hold a thin tubule of membrane at a steady-state length (Dai and Sheetz, 1995; Sheetz, 2001). Although tether forces are measured by externally imposed mechanical perturbations, they offer an approximation of the mechanical challenge that the cytoskeletal machinery faces during protrusion formation and suggest that pN-scale pushing force is needed. A long history of previous theoretical and experimental studies have argued that actin polymerization generates this force during protrusion growth (Condeelis, 1993; Hill and Kirschner, 1982a; Pollard and Borisy, 2003). Indeed, the incorporation of new actin subunits into the growing barbed ends of actin filaments generates force in the required pN-scale range (Kovar and Pollard, 2004). Additionally, because protrusions are supported by 10s to 100s of parallel filament bundles, these structures should be able to generate outward pushing force that exceeds the threshold suggested by tether force estimates. However, the ability of a growing actin filament to generate force also depends on the availability of free actin monomers (Theriot, 2000). This poses a mechanistic conundrum given the tight confines of the intrafilopodial cytoplasm, where the diffusive availability of new monomers at growing barbed ends could become limiting as a protrusion elongates. From this perspective, we set out to test the hypothesis that myosin motors, which are abundant residents of varying actin-based protrusions, provide an additional source of mechanical force for driving protrusion growth.

Based on their mechanical potential and localization at the interface between polymerizing actin filaments and cellular membranes, myosins are well-positioned to drive membrane deformation in diverse biological contexts. Driving the growth of surface membrane protrusions is a long-standing functional hypothesis proposed for several unconventional members of the myosin superfamily, including motors from classes 1, 3, 5, 7, 10, and 15 (Houdusse and Titus, 2021). Previous cell biological studies have emphasized cargo transport models, which propose that protrusion myosins deliver specific factors to the distal tip compartment, and these in turn promote growth via a range of molecular mechanisms. We now know that myosins in microvilli, filopodia, and stereocilia are critical for the distal tip enrichment of scaffolding proteins (Crawley et al., 2014; Crawley et al., 2016; Grati and Kachar, 2011; Li et al., 2017), adhesion molecules (Weck et al., 2016; Yu et al., 2017), and proteins that directly impact actin dynamics and organization (Arthur et al., 2021; Ebrahim et al., 2016; Manor et al., 2011; Tokuo and Ikebe, 2004). However, understanding a given myosin’s contributions to protrusion growth, in terms of tip-directed cargo delivery vs. more direct mechanical effects, has proved challenging largely because an experimental approach for isolating the impact of motor domain force generation in cells has remained elusive.

In this report, we describe a novel cell-based system that enabled us to directly test whether myosin generated force contributes to protrusion elongation. Using Myo10 as a model myosin, we found that—independent of any cargo binding potential—Myo10 motor domains drive robust filopodial growth when docked onto the plasma membrane using the rapalog-based system described here. Leveraging this approach to develop further insight into the properties of induced filopodial growth, we learned that elongation requires active motor domains that exert barbed end-directed force. Moreover, the elongation of individual protrusions was paralleled by an accumulation of Myo10 motor domains at filopodial tips, strongly suggesting that the local level of force application to the membrane controls filopodial growth kinetics. Importantly, induced filopodial elongation was supported by structurally distinct myosin motors, and membrane binding modules that associate with the plasma membrane using different mechanisms (transmembrane vs. peripheral). We, therefore, propose that the motor-driven filopodial growth revealed in our studies likely reflects a general activity of different unconventional myosins in diverse actin-based protrusions.

Why does docking myosin motor domains onto the plasma membrane drive such robust filopodial growth? We propose that motor domains impart barbed end-directed force directly onto the membrane that encapsulates a growing filopodium, and this, in turn, reduces the physical barrier to elongation. One possibility is that myosin-generated force “softens” the membrane (i.e., lowers membrane tension) in the distal tip compartment, allowing actin monomers to incorporate into the barbed ends of core bundle filaments more readily. Consistent with this idea, we found that acute expansion of plasma membrane with the addition of a lipid intercalating probe, immediately led to additional filopodial elongation. This concept is also supported by biophysical studies showing that application of external force to the tip of a filopodium accelerates actin incorporation at the distal end (Bornschlogl and Bassereau, 2013). A second possibility is that force generated by membrane-bound motor domains contributes directly to the retrograde displacement of the actin core bundle; this again would be expected to ease the barrier to actin monomer incorporation at the distal ends. This latter idea finds support in previous studies on actin treadmilling dynamics in neuronal growth cone filopodia and epithelial microvilli (Chinowsky et al., 2020; Medeiros et al., 2006). A more likely scenario is that myosin-derived force exerts some combination of both effects.

One open question that re-emerges here is whether a myosin motor can impart mechanical force to a fluid lipid bilayer. *In vitro* studies have shown that membrane lipids are generally free to diffuse through the plane of the bilayer, as indicated by relatively high diffusion coefficients (Sanderson, 2012). Based on this, one might expect that mechanical force imparted to proteins in the bilayer (e.g. CDHR2TM) or attached to the surface of the bilayer (e.g. TH1, BTK) would dissipate quickly. However, experimental evidence from more complex lipid mixtures that include cholesterol, indicate much lower rates of diffusion with significant frictional coefficients (Cicuta et al., 2007), leading to the possibility that external force applied by myosin motors may persist for a period of time. To be matched to the kinetics of motor domain attachment to actin, this duration only needs to be in the range of milliseconds (e.g. Myo10 exhibits a strongly bound duration of ∼80 ms)(Takagi et al., 2014). Moreover, filopodia contain 100s to 1000s of Myo10 molecules, and individual motors will cycle asynchronously and, thus, apply force continuously to the overlying plasma membrane. We, therefore, expect that protrusion resident myosins can exert biologically significant (pN scale) levels of force on the membrane for prolonged periods of time.

Of the numerous unconventional myosin motor domains tested for their ability to support filopodial elongation in this work, only representatives from classes 3, 10, and 15 were able to drive growth. Do these myosins share common features that explain this selectivity? One possible explanation might be linked to the biochemical composition of the core actin filament bundles that support filopodia. Fascin-1 is the dominant bundling protein in these structures and *in vitro* studies have revealed that the filopodial myosin, Myo10, preferentially interacts with and moves processively along actin filaments that are crosslinked in parallel by this bundler (Ropars et al., 2016; Takagi et al., 2014). Interestingly, as residents of stereocilia, Myo3 and Myo15 are adapted for interacting with core bundles organized by fascin-2 in addition to espin and plastin (Krey et al., 2016; Perrin et al., 2013; Sekerkova et al., 2006). Microvilli are structurally distinct from both filopodia and stereocilia in that their filaments are bundled by plastin, villin, espin and MISP (Bartles et al., 1998; Bretscher and Weber, 1979, 1980; Morales et al., 2022), but not fascin. Thus, a preference for the close filament spacing that is characteristic of fascin bundling might explain the robust filopodial elongation response demonstrated by the motor domains of Myo3, Myo10, and Myo15, relative to other myosins tested in our assay. Another possible explanation for why only a subset of myosins can induce filopodial elongation may be related to the natural diversity of kinetic properties exhibited by different motors. Indeed, myosins from different superfamily classes exhibit a broad range of actin filament sliding speeds and ATP hydrolysis kinetics (O’Connell et al., 2007). It follows that certain kinetic properties might be needed to support robust filopodial elongation in the assay we describe here. One might expect that, at a minimum, any motor capable of driving filopodial growth would need to demonstrate a barbed end-directed mechanical velocity faster than the actin retrograde flow rate. Velocities measured from sliding filaments assays or direct single-molecule observations indicate speeds for Myo3a, Myo10, and Myo15 in the range of 100s of nm/sec (110 nm/s, 310-660 nm/s, 278-429 nm/s, respectively)(Bird et al., 2014; Komaba et al., 2003; Ropars et al., 2016; Takagi et al., 2014), approximately 10-fold higher than typical retrograde flow rates (Bornschlogl et al., 2013). Thus, at least in the case of these three motors, this requirement appears to be met. Another kinetic property relevant here is the duty ratio: the fraction of the total ATPase cycle time that a motor domain will remain bound to actin. A high duty ratio myosin will be able to exert higher time-average force, which will increase its mechanical impact when present in a multi-motor ensemble (Bird et al., 2014; Dose et al., 2007; Homma and Ikebe, 2005).

Unconventional myosin isoforms have long been recognized as abundant components of actin-based protrusions. Whether these motors contribute growth-promoting force has remained unclear, primarily because of the technical challenges associated with uncoupling the roles of cargo transport from more direct physical effects. The data we report here indicates that myosins are also able to contribute significant growth-promoting force to drive robust protrusion elongation. As one of the main limitations in studying surface protrusions is the stochastic nature of their assembly, we expect that the synthetic, inducible system introduced in this study will prove useful for biologists seeking to uncover a deeper understanding of the biochemical and biophysical nature of protrusion formation.

## ACKNOWLEDGMENTS

The authors would like to thank all members of the M.J.T. laboratory, members of the Vanderbilt Microtubules and Motors Club, and L.M.M. for their feedback and guidance. Light microscopy was performed in part through use of the VUMC Cell Imaging Shared Resource. This work was supported by the NIH NIDDK National Research Service Award F31DK130599 (G.N.F.), and NIH grants R01-DK125546, R01-DK095811, and R01-DK111949 (M.J.T.).

## AUTHOR CONTRIBUTIONS: G.N.F. M.W. C.B. O.L.P. M.J.T

Conceptualization, G.N.F. and M.J.T.; Methodology, G.N.F., M.W., M.J.T.; Software, O.L.P.; Validation, G.N.F. and C.B.; Formal Analysis, G.N.F. and O.L.P.; Investigation, G.N.F. and C.B.; Writing-Original Draft, G.N.F. and M.J.T.; Writing-Review & Editing, all authors; Visualization, G.N.F.; Supervision, M.J.T.; Project; Administration, M.J.T.; Funding Acquisition, G.N.F. and M.J.T.; All authors contributed to revising the manuscript.

## SUPPLEMENTAL FIGURE LEGENDS

**Figure S1:**
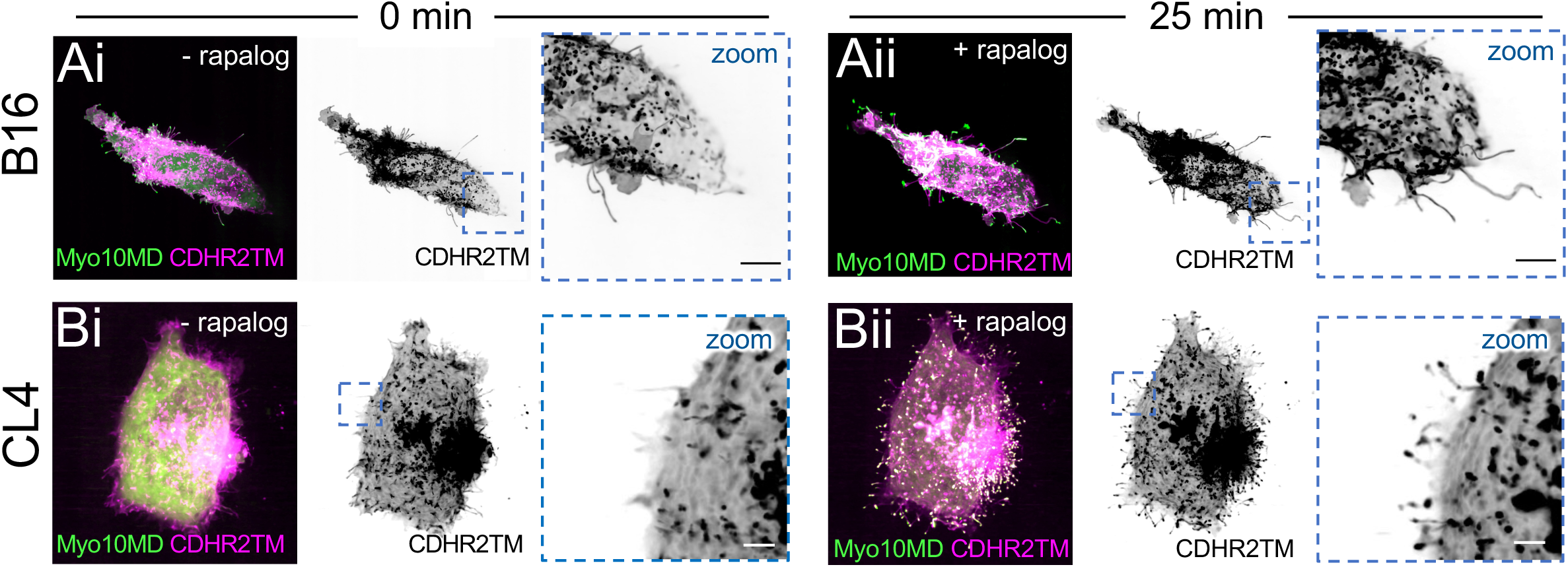
Filopodia induced by membrane bound Myo10MD is functional in distinct cell types. (**A**) Merged confocal maximum intensity projection of B16 cells expressing Myo10MD (green) and CDHR2TM (magenta) at 0 min (**Ai**) and 25 min after (**Aii**) rapalog treatment. (**B**) Merged confocal maximum intensity projection of CL4 cells expressing the Myo10MD (green) and CDHR2TM (magenta) at 0 min (**Ai**) and 25 min after (**Aii**) rapalog treatment. Scale bars = 10 μm.

**Figure S2.**
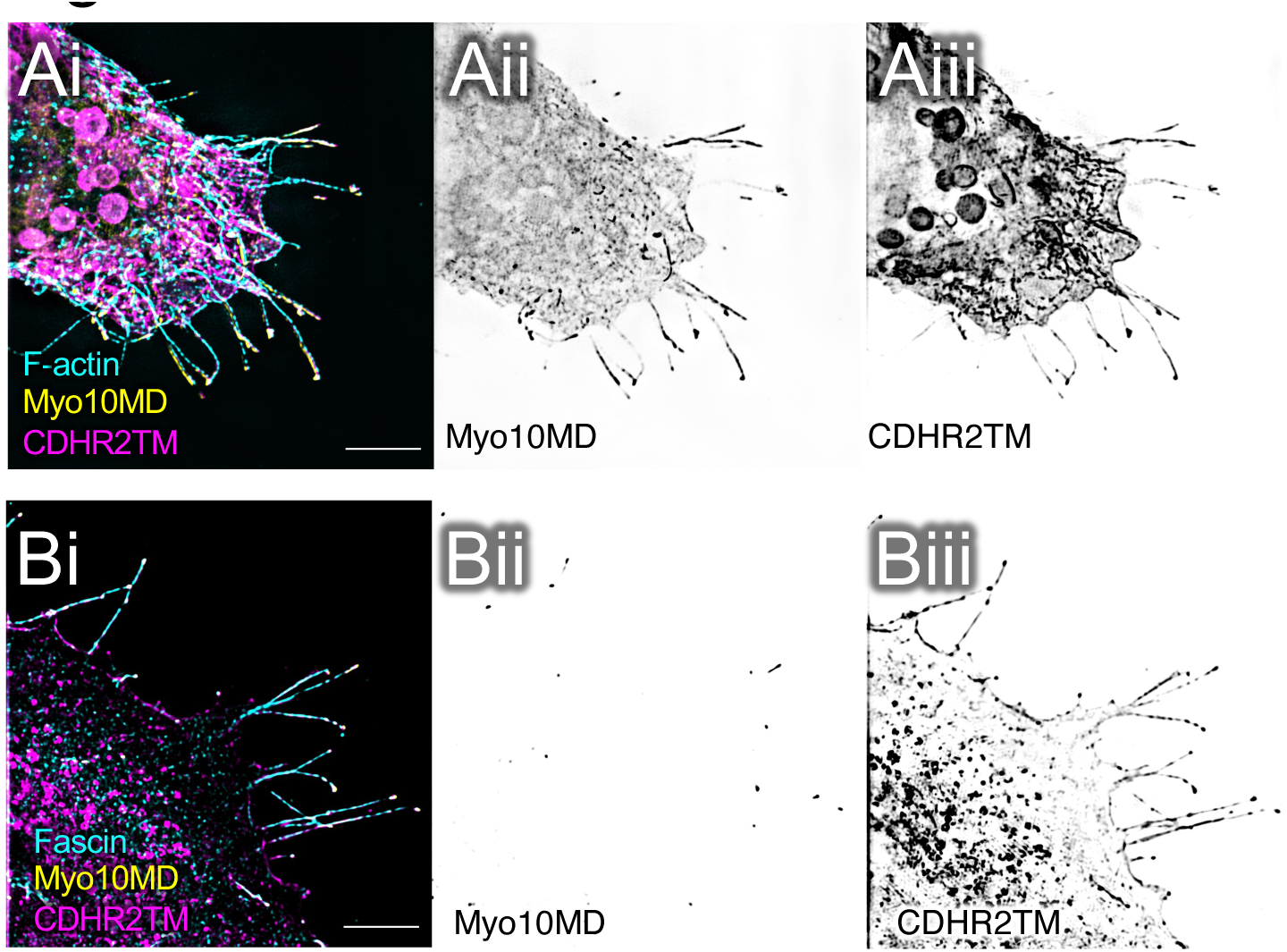
(accompanies Fig 2A and 2B): Expression of the membrane bound Myo10MD system. Inverted single channel SIM image of Myo10MD (Aii and Bii) and CDHR2TM (Aiii and Biii) to confirm expression of these constructs shown in Fig 2A and 2B. Scale bar = 5 μm.

**Figure S3:**
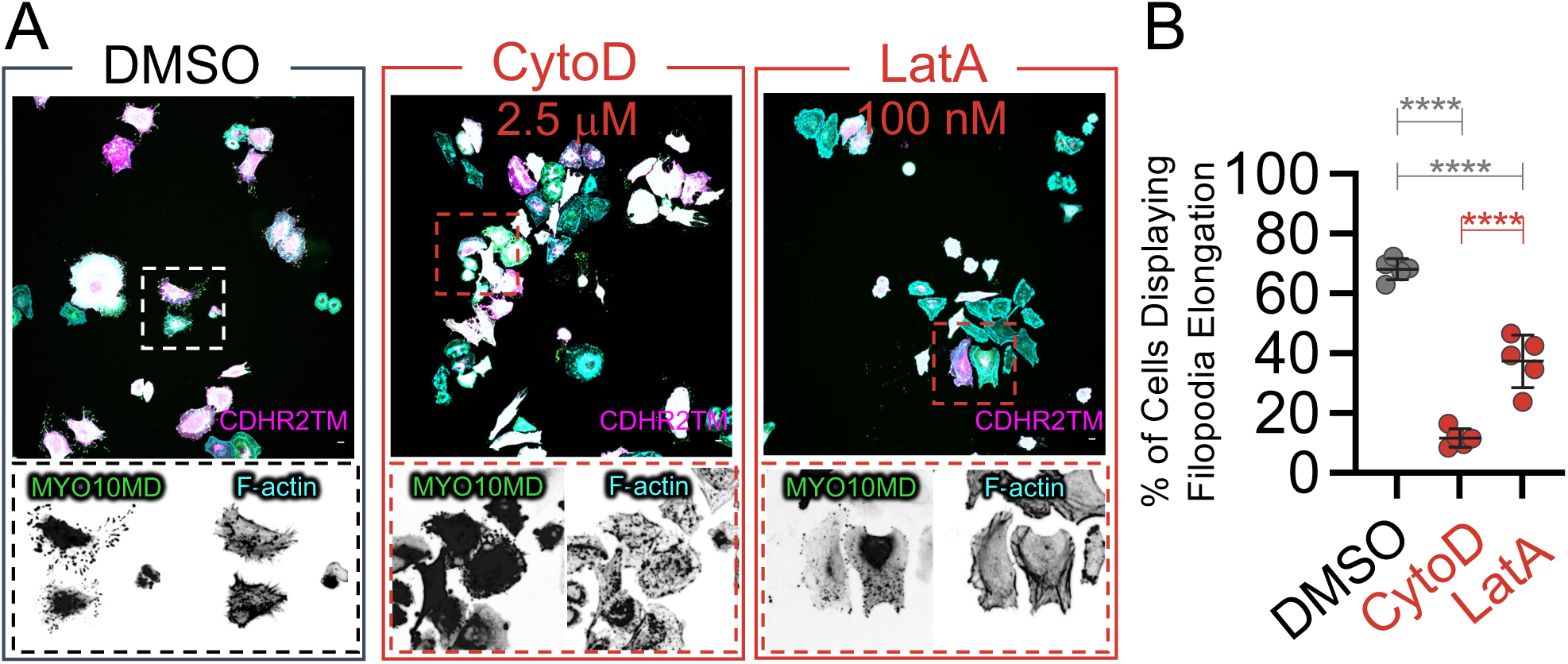
Filopodia induced by membrane bound Myo10MD requires polymerizing actin. (A) Merged confocal maximum intensity projection of HeLa cells expressing the Myo10MD (green) and CDHR2TM (magenta) constructs 25 min after rapalog treatment and either DMSO (control), 2.5 μm Cytochalasin D (CytoD), or 100 nM Latrunculin A (LatA). Scale bar = 10 μm. (B) Quantification of the percentage of cells displaying filopodia growth after treatment with rapalog and either DMSO, CytoD, or LatA; n = > 370 cells for each treatment; error bars represent the mean ± SD; ****p-value ≤ 0.0001 was calculated using an Ordinary one-way ANOVA followed by Tukey’s multiple comparisons test to compare treatments with each other.

**Figure S4:**
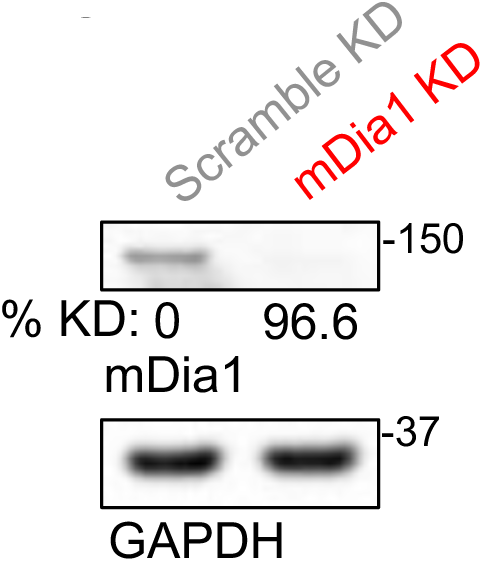
Expression of DIAPH1 (mDia1) in knockdown (KD) cells. (A) Representative western blot analysis of expression levels of mDia1 in lysates from HeLa cells siRNA scramble control (Scramble KD) and siRNA mDia1 KD (mDia1 KD). GAPDH was used as a loading control. Percent KD below mDia1 expression blot based on densitometry measurements normalized to GAPDH.

**Figure S5:**
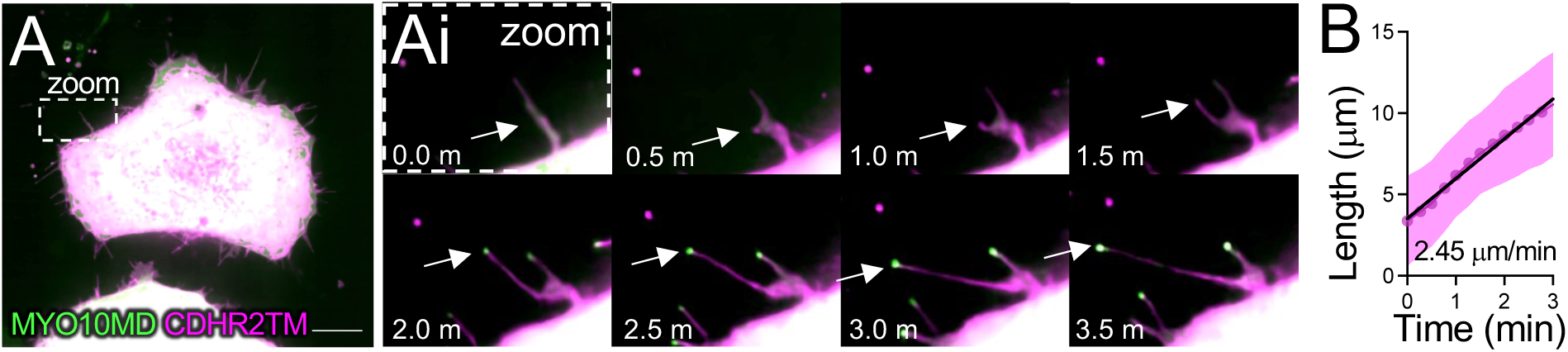
Methylcellulose does not alter the growth rates of filopodia induced by membrane bound Myo10MD. (**A**) Merged confocal maximum intensity projection of HeLa cell expressing Myo10MD (green) and CDHR2TM (magenta) after the addition of rapalog. Scale bar = 10 μm. (**Ai**) Montage of zoomed in region (**A**) highlighting the growth of a single filopodium (white arrow). (**D**) Quantification of the elongation rate of single filopodium induced by rapalog treatment in media containing 0.5% methylcellulose. Rate was calculated via the slope using a simple linear regression; *n* = 10 individual filopodia from n = 2 separate cells. Error bars represent the SD.

**Figure S6:**
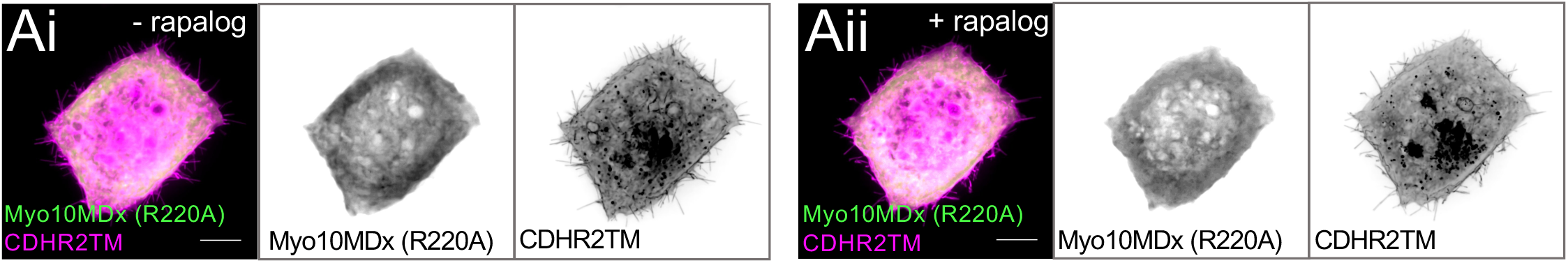
Myosin-10 motor domain switch I dead (R220A) construct does not elongate filopodia. Merged confocal maximum intensity projection of HeLa expressing Myo10MD-R220A (green) and CDHR2TM (magenta) at 0 min (**Ai**) and 25 min after (**Aii**) rapalog treatment. Inverted single channel images shown to the right of the merged. Scale bar = 10 μm.

**Figure S7:**
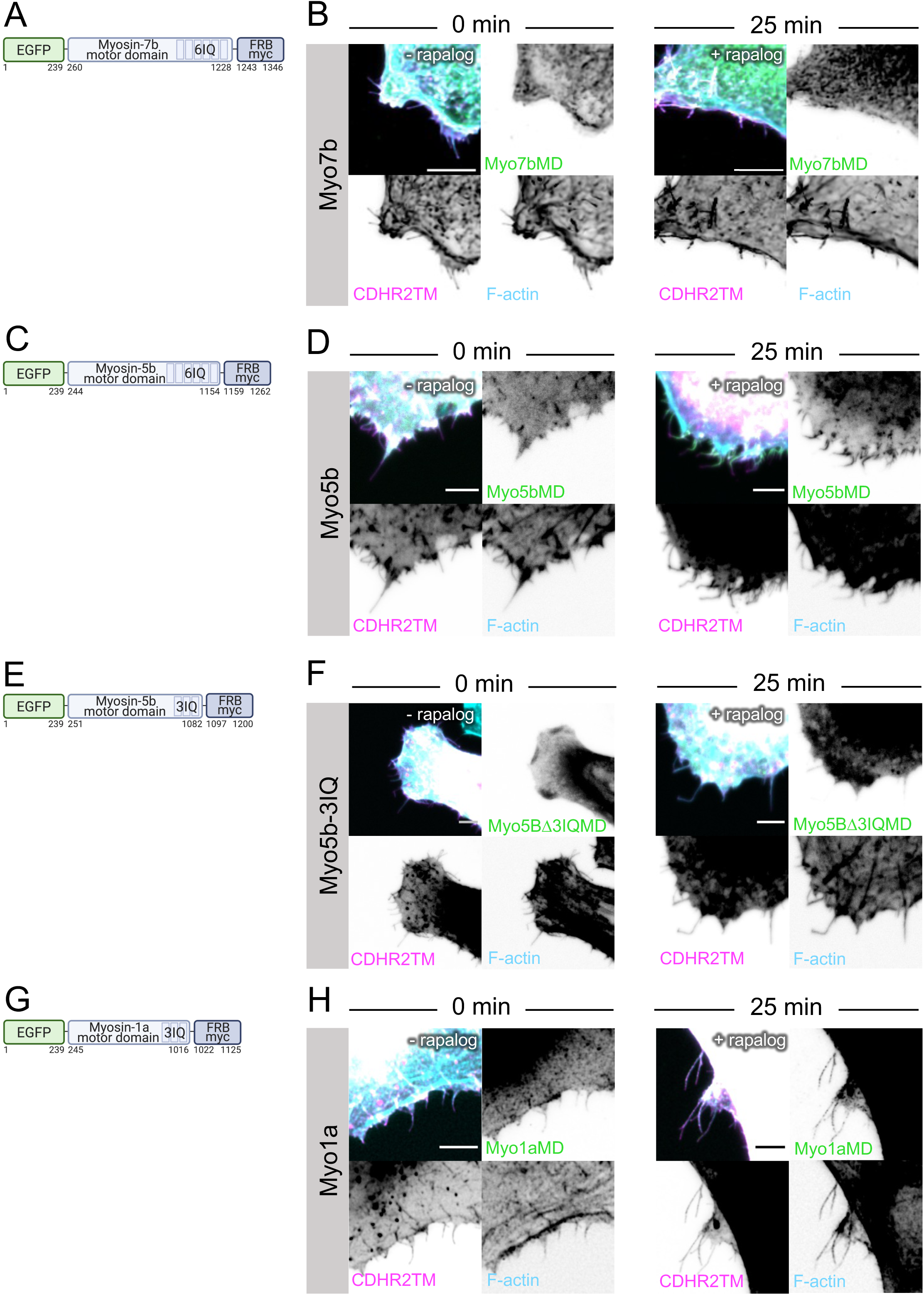
Myosin-7b, -5b, -5bΔ3IQ and -1a motor domains do not elongate filopodia. (**A**) Cartoon diagram showing the EGFP-Myosin-7b Motor Domain (Myo7bMD)-FRB-myc construct. (**B**) Merged confocal maximum intensity projection of HeLa cells expressing Myo7bMD (green) and CDHR2TM (magenta) at 0 min and 25 min after rapalog treatment. (**C**) Cartoon diagram showing the EGFP-Myosin-5b Motor Domain (Myo5bMD)-FRB-myc construct. (**D**) Merged confocal maximum intensity projection of HeLa cells expressing Myo5bMD (green) and CDHR2TM (magenta) at 0 min and 25 min after rapalog treatment. (**E**) Cartoon diagram showing the EGFP-Myosin-5b Motor Domain with 3 IQ domains deleted (Myo5bdel3IQMD)-FRB-myc construct. (**F**) Merged confocal maximum intensity projection of HeLa cells expressing Myo5bdel3IQMD (green) and CDHR2TM (magenta) at 0 min and 25 min after rapalog treatment. (**G**) Cartoon diagram showing the EGFP-Myosin-1a Motor Domain (Myo1aMD)-FRB-myc construct. (**H**) Merged confocal maximum intensity projection of HeLa cells expressing Myo1aMD (green) and CDHR2TM (magenta) at 0 min and 25 min after rapalog treatment. Inverted single channel images shown to the right and below of the merged. All images were also stained for F-actin (cyan). Scale bar = 5 μm.

## SUPPLEMENTAL VIDEO LEGENDS

**Video S1.** Myo10MD no rapalog. Live cell movie of maximum intensity projections of HeLa cell expressing Myo10MD-FRB (green) and CDHR2TM-FKBP (magenta) not treated with rapalog. 3 μm maximum intensity projections are composed of 16 x 0.2 μm confocal slices. Images were captured at 1 frame per 5 min. Time min:sec.

**Video S2.** Myo10MD rapalog treatment. Live cell movie of maximum intensity projections of HeLa cell expressing Myo10MD-FRB (green) and CDHR2TM-FKBP (magenta) treated with rapalog. 3 μm maximum intensity projections are composed of 21 x 0.15 μm confocal slices. Images were captured at 1 frame per 15 sec. Time min:sec.

**Video S3.** Single filopodium growth. Live cell movie of maximum intensity projections of HeLa cell expressing Myo10MD-FRB (green) and CDHR2TM-FKBP (magenta) treated with rapalog. 3 μm maximum intensity projections are composed of 3 x 0.2 μm confocal slices. Images were captured at 1 frame per ∼2 sec. Time min:sec. Scale bar = 5 μm.

**Video S4**. Myo10MDx. Live cell movie of maximum intensity projections of HeLa cell expressing Myo10MDx-FRB (green) and CDHR2TM-FKBP (magenta) treated with rapalog. 3 μm maximum intensity projections are composed of 17 x 0.2 μm confocal slices. Images were captured at 1 frame per 15 sec. Time min:sec.

**Video S5.** Myo6MD. Live cell movie of maximum intensity projections of HeLa cell expressing Myo6MD-FRB (green) and CDHR2TM-FKBP (magenta) treated with rapalog. 3 μm maximum intensity projections are composed of 11 x 0.3 μm confocal slices. Images were captured at 1 frame per 15 sec. Time min:sec.

**Video S6.** Lat A CTRL. Live cell movie of maximum intensity projections of HeLa cell expressing Myo10MD-FRB (green) and CDHR2TM-FKBP (magenta) treated with rapalog after a 7-day incubation with Latrunculin A (LatA) for 75 min. 3 μm maximum intensity projections are composed of 16 x 0.2 μm confocal slices.

**Video S7.** Lat A washout. Live cell movie of maximum intensity projections of HeLa cell expressing Myo10MD-FRB (green) and CDHR2TM-FKBP (magenta) treated with rapalog after a 7-day incubation with Latrunculin A (LatA) and then with a LatA washout. Cells were imaged for 75 min with the Lat A washout at 35 min. 3 μm maximum intensity projections are composed of 18 x 0.18 μm confocal slices. Images were captured at 1 frame per 5 min. Time min:sec.

**Video S8.** CellMask CTRL. Live cell movie of maximum intensity projections of HeLa cell expressing Myo10MD-FRB (green) and CDHR2TM-FKBP (magenta) treated with rapalog for 140 min. 3 μm maximum intensity projections are composed of 18 x 0.18 μm confocal slices. Time hr:min:sec.

**Video S9.** CellMask addition. Live cell movie of maximum intensity projections of HeLa cell expressing Myo10MD-FRB (green) and CDHR2TM-FKBP (magenta) treated with rapalog after a 7-day incubation with Latrunculin A (LatA) and then with a LatA washout at 35 min and imaged for 140 min. 3 μm maximum intensity projections are composed of 18x 0.18 μm confocal slices. Images were captured at 1 frame per 5 min. Time hr:min:sec.

**Video S10.** Myo3aΔKMD. Live cell movie of maximum intensity projections of HeLa cell expressing Myo3aΔKMD-FRB (green) and CDHR2TM-FKBP (magenta) treated with rapalog. 3 μm maximum intensity projections are composed of 16 x 0.2 μm confocal slices. Images were captured at 1 frame per 5 min. Time min:sec.

**Video S11.** Myo15aMD. Live cell movie of maximum intensity projections of HeLa cell expressing Myo15aMD-FRB (green) and CDHR2TM-FKBP (magenta) treated with rapalog. 3 μm maximum intensity projections are composed of 18 x 0.18 μm confocal slices. Images were captured at 1 frame per 5 min. Time min:sec.

## STAR METHODS

### Plasmids

All PCR amplification was completed using Q5 High-Fidelity DNA Polymerase (NEB #M0491S) unless site-mutagenesis was completed, in which case PfuUltra High-Fidelity DNA Polymerase (in Agilent #200523 kit) was used.

#### Membrane interacting motifs

To generate the membrane binding motif constructs, we first generated a (1) <u>mCherry-FKBP-myc</u> plasmid by inserting mCherry (a.a. 1329-1564) from CDHR2-mCherry-FKBP-myc using the primers 5’- taagcagaattccatggtgagcaagggcga-3’ and 5’-taagcagcggccgccactgtg-3’ to amplify a PCR product of mCherry flanked by the restriction digest sites for EcoRI (NEB #R3101) and NotI (NEB #R3189). The gel purified mCherry PCR product and pcDNA3.1 Zeo+FKBP-myc (Addgene plasmid #20211) were digested with EcoRI and NotI, gel purified (Macherey-Nagel #740609.50), and ligated together (NEB M2200S). (3) <u>CDHR2TM-</u> <u>mCherry-FKBP-myc</u> was made by amplifying the single-spanning transmembrane domain of a previously generated full-length CDHR2-mCherry-FKBP-myc plasmid. The transmembrane portion of the protein (a.a. 1-185) was amplified using the primers 5’- taagcagctagcaccatggcccagctatgg-3’ and 5’-taagcacttaagcaggtccgtggtgtccagg-3’ to generate a PCR product flanked by the restriction sites NheI (NEB #R0131) and AflII (NEB #R0520). Using traditional cloning, this fragment was then inserted into the mCherry-FKBP-myc plasmid. To generate the (2) <u>mCherry-BTK-PH-FKBP-myc</u> PI(3,4,5)P_3_ membrane binding motif plasmid, we amplified the BTK-PH domain of *hs* Bruton tyrosine kinase (NM_000061.2) from Addgene plasmid #51463 (a.a. 1-177) using the primers 5’-tgcttagaattcggttttaagcttccattcctgttctcca-3’ and 5’-taagcaggtaccaccatggccgcagtgattctggag-3’ to generate a PCR product containing BTK-PH flanked by the restriction digest sites KpnI and EcoRI. The mCherry-FKBP-myc plasmid and gel purified BTK-PH PCR product were both digested with KpnI (NEB #R3142) and EcoRI, gel purified and ligated together. To generate the (3) <u>FKBP-myc-</u> <u>mCherry-TH1</u> electrostatic membrane binding motif plasmid, Gibson assembly (NEB #E2621) was used to assemble three fragments into pcDNA3.1 Zeo+FKBP-myc: Fragment 1: FKBP-myc (a.a. 1-117) from pcDNA3.1 Zeo FKBP-myc; Fragment 2: mCherry (a.a. 1-236) from the previously generated full length CDHR2-mcherry-FKBP-myc ; Fragment 3: TH1 domain (a.a. 772-1043) of *hs* unconventional myosin-1a (GeneBank^TM^ AF009961) was amplified using the primers 5’- ggcggccgctcgagtctagagccctcaccttggcaga-3’ and 5’- gaaggcacagtcgaggctgatcagcgggtttaaacgggcctcactgcacagtcacctccaag-3’.

#### Myosin motor domains

We first generated the (1) <u>EGFP-Myo10MD-FRB-myc</u> construct by inserting the motor domain of *Hs* myosin-10 (a.a. 1-855) into the pcDNA3.1 Zeo+FRB-myc (Addgene plasmid #20228) plasmid by Gibson assembly. The motor domain was amplified was using the primers 5’- tatagggagacccaagctgggccaccatggtgagcaag-3’ and 5’- actgtgctggatatctgcagcctgcttcctccgttgggc-3’ and the FRB-myc plasmid was linearized by restriction enzyme incubation with NheI and EcoRI. For the additional motor domain constructs, we generated an (2) <u>EGFP-FRB-myc</u> plasmid to insert various motor domains into by amplifying EGFP (a.a. 1-239) from EGFP-Myo10MD-FRB using the primers 5’- taagcaaagcttaccatggtgagcaagggcg-3’ and 5’-taagcagaattcagatctgagtccggacttgtacagc-3’ to generate a PCR product containg EGFP flanked by HindIII and EcoRI. The FRB-myc plasmid and gel purified EGFP PCR product were both digested with HindIII (NEB #R3104) and EcoRI, gel purified and ligated together. To generate the two motor domain dead constructs (G437A, switch II and R220A, switch I) we utilized site-directed mutagenesis (Agilent #200523) of EGFP-Myo10MD-FRB to induce the two, separate amino acid changes using the primers 5’-cacctcaaagttttcaaatgcaaagatgtccaagatgcc-3’ and 5’-ggcatcttggacatctttgcatttgaaaactttgaggtg-3’ for (3) <u>EGFP-Myo10MDxG437A-FRB-</u> <u>myc</u> and 5’-ctgaacaaacttcccgaaggcactggagttattgttgtac-3’ and 5’- gtacaacaataactccagtgccttcgggaagtttgttcag-3’ for (4) <u>EGFP-Myo10MDxR220A-FRB-myc</u>. To generate the (5) <u>EGFP-Myo6MD-FRB-myc</u> plasmid, Gibson assembly was used to assemble three fragments of *hs* unconventional myosin-6 (Genebank #NM_001300899.2) to replace the Myo10MD in EGFP-Myo10MD-FRB-myc: Fragment 1: a.a. 1-294 was amplified using the primers 5’-agatctcgaatcacaagtttgtacaaaaaagcaggctccgaaatggaggatggaaagcccgt-3’ and 5’- cggttctgtaaaatctgtttgtcagtttcttt-3’; Fragment 2: a.a. 295-576 was amplified using the primers 5’-ttgctaacaaagaaactgacaaacagattttacag-3’ and 5’- gccttcgtcgtctctgatattcctatgaactg-3’; Fragment 3: a.a. 577-844 was amplified using the primers 5’-taagctggcagttcataggaatatcagagacg-3’ and 5’- cagaggattctagactcgagcggccgccactgtgctggattgcccaccttaaccagacca-3’. The backbone (EGFP-Myo10MD-FRB-myc) was linearized by restriction digest via EcoRV (NEB #R3195) and EcoRI. The myosin-3a lacking the N-terminal kinase (ΔK) plasmid was kindly gifted to us by Dr. Christopher Yengo. (6) <u>EGFP-Myo3aΔK-FRB-myc</u> was generated using Gibson assembly to insert the Myo3aΔKMD (a.a. 407-1432) into the EGFP-FRB-myc plasmid. The Myo3aΔMD was amplified using the primers 5’- cacagtggcggccgctcgaggtagatgatttagcaaccctagaagttttgga-3’ and 5’- cacatctcatgccagaggataaaagaagacatcttgtccttcctcaact-3’, and the EGFP-FRB-myc plasmid was linearized using the primers 5’-aggacaagatgtcttcttttatcctctggcatgagatgtgg-3’ and 5’-agggttgctaaatcatctacctcgagcggccgcc-3’. The *Mm* myosin-15a (NM_182698) plasmid was kindly gifted to us by Dr. Uri Manor. The (7) <u>EGFP-Myo15aMD-FRB-myc</u> plasmid was generated using Gibson assembly to insert the motor domain of myo15a (a.a. 1-750) into EGFP-FRB-myc. The Myo15aMD was amplified using the primers 5’- cacagtggcggccgctcgagatgcactccatacgcaacctgcc-3’ and 5’- atctcatgccagaggattctcagcagactctgcctcatct-3’, and the EGFP-FRB-myc backbone was linearized by PCR using the primers 5’- agatgaggcagagtctgctgagaatcctctggcatgagatgtggc-3’ and 5’- aggttgcgtatggagtgcatctcgagcggccgcc-3’. The (8) <u>EGFP-Myo7bMD-FRB-myc</u> plasmid was generated using Gibson assembly to insert EGFP and the motor domain (a.a. 1 - 968) of *Hs* myosin-7b (NM001393586.1) into FRB-myc. EGFP and the myosin-7b motor domain were amplified by PCR using the primers 5’- tatagggagacccaagctgggccaccatggtgagcaag-3’ and 5’- actgtgctggatatctgcagcccaggccatccacatcctc-3’ and the FRB-myc backbone was linearized by restriction digest with the enzymes NheI and EcoRI. The *Hs* myosin-5b plasmid was kindly gifted to us by Dr. James R. Goldenring. The (9) <u>EGFP-Myo5bMD-FRB-myc</u> plasmid was generated using Gibson assembly to insert the motor domain of myosin-5b (a.a. 1-910) into EGFP-FRB-myc. The myosin-5b motor domain fragment was amplified using the primers 5’-Acaagtccggactcagatctatgagcgtgggcgagc-3’ and 5’- aggattctagactcgagcggggcctcgattctcagggc-3’, and the EGFP-FRB-myc plasmid was linearized by PCR using the primers 5’-aagccctgagaatcgaggccccgctcgagtctagaatcctctgg-3’ and 5’-tacagctcgcccacgctcatagatctgagtccggacttgtacag-3’. The (10) <u>EGFP-</u> <u>Myo5bΔ3IQ-FRB-myc</u> was generated by amplifying a fragment containing the motor domain and first three IQ motifs (a.a. 1-831) from EGFP-Myo5b-FRB-myc, flanked by EcoRI sites using the primers 5’-taagcagaattctccggactcagatctatgagcgtgg-3’ and 5’- taagcagaattcggcctgtctggcccg-3’. Using traditional cloning, this fragment was then inserted into EGFP-FRB-myc. The (11) <u>EGFP-Myo1aMD-FRB-myc</u> plasmid was generated by amplifying the motor domain (a.a. 1-771) of *Hs* myosin-1a (NM001256041.2) using the primers 5’-agtccggactcagatctgaaatgcctctcctggaaggt-3’ and 5’-attctagactcgagcggccgagcctctgaccggaaatatttgc-3’. Using Gibson assembly, this fragment was then inserted into a PCR linearized EGFP-FRB-myc backbone using the primers 5’-aatatttccggtcagaggctcggccgctcgagtctaga-3’ and 5’gaaccttccaggagaggcatttcagatctgagtccggacttgtaca-3’. For all plasmids, colonies were first screen by restriction digest to confirm the correct digested band pattern and then verified by sequencing (GENEWIZ, South Plainfied, NJ).

### Cell culture

HeLa, HEK293FT, B16-F1, and LLC-PK1-CL4 (CL4), cells were cultured at 37°C and 5% CO_2_ in Dulbecco’s modified Eagle’s medium (DMEM) (Corning #10-013-CV) with high glucose and 2 mM L-Glutamine supplemented with 10% fetal bovine serum (FBS). Transfections were performed using Lipofectamine 2000 (Thermo Fischer #11668019) according to the manufacturer’s protocol. Cells were incubated in Lipofectamine for 4-6 hrs, after which they were replated onto 35mm glass-bottom dishes (Cellvis #D35-20-1.5-N) and/or coverslips coated with 25 ug/mL laminin (Corning #354232) and allowed to adhere and recover overnight before live imaging or fixing/staining. For DIAPH1 (mDia1) knock down (KD) experiments, HeLa cells were transfected twice with either 10 μM of Accell human DIAPH1 siRNA SMARTPool (Horizon Discovery #E-010347-00-0005) or Accell non-targeting control pool (Horizon Discovery #D-001910-10-05). Cells were allowed to recover overnight before being transfected a second time with their respevtive siRNA pools. On the third day, HeLa cells were transfected with the EGFP-Myo10MD-FRB and CDHR2TM-mCherry-FKBP constructs and replated as described above.

### Immunofluorescence

For Structured Illumination Microscopy (SIM) and A1 confocal imaging, cells were washed with pre-warmed phosphate-buffered saline (PBS) before being fixed in 4% paraformaldehyde (Electron Microscopy Sciences #15710) in PBS for 15 mins at 37°C. After fixation, cells were rinsed three times in PBS and permeabilized with 0.1% Triton X-100 in PBS for 15 mins at 37°C. If cells were stained with anti-Fascin, cells were fixed and permeabilized with ice cold methanol on ice for 15 mins instead of 4% paraformaldehyde. After permeabilization, cells were rinsed three times in PBS and blocked with 5% BSA (Research Products International #9048-46-8) in PBS for 1 hr at 37°C. Immunostaining was performed using the primary antibodies: anti-VASP (1:50; Cell Signaling Technologies #3132S), anti-Fascin (1:100; Agilent Technologies #M356701-8), anti-GFP (1:200; Aves Labs #GFP-1020), anti-mCherry (1:500; Invitrogen #M11217), diluted in 1% BSA in PBS at 37°C for 1 hr. After incubation with primary antibody, coverslips were rinsed three times in PBS and incubated for 1 hr with Alex Fluor 568-phalloidin or Alex Fluor 647-phalloidin (1:200; Invitrogen #A12380 and #A22287) at room temperature. Coverslips were then washed five times in PBS and mounted on glass slides in ProLong Gold (Invitrogen #P36930).

### Drug Treatment

To oligomerize FRB and FKBP constructs, transfected cells were treated with 500 nM of A/C Heterodimerizer (Takara #635057) for 25 mins before being fixed for immunofluorescence or were treated with 500 nM of A/C Heterodimerizer at the onset of live imagining. To inhibit actin polymerization, HeLa cells were treated with either 2.5 uM Cytochalasin D (Sigma #C2618) or 100 nM Latrunculin A (Lat A; Molecular Probes #12370) in addition to 500 nM of the A/C Heterodimerizer, and fixed after 25 mins. For the Lat A washout experiments, cells were treated with 100 nM Lat A. For the live-cell imaging membrane addition experiments, cells were treated with CellMask^TM^ (Thermo #C10045) at 1:1000. For live-cell imaging of individual filopodia, 0.5% methylcellulose diluted in DMEM with high glucose and 2 mM L-Glutamine supplemented with 10% FBS was added prior to imaging.

### Western Blot Analysis of Cultured Cells

To generate lysates for western blotting to confirm DIAPH1 (mDia1) KD, HeLa cells were cultured in 6-well plates, and cell lysates were prepared on ice using Cellytic M lysis buffer (Sigma #C2978) supplemented with 4mM pefabloc (sc-202041), a pellet protease inhibitor (Roche #04906845001) and 2mM ATP (Sigma #A1852). Samples were centrifuged at 12,000 rpm for 10 min at 4°C to remove cell debris. The resulting supernatant was normalized and boiled in Laemmli Sample Buffer (Bio-Rad #1610737) with 5% beta-mercaptoethanol (Sigma #M3148) for 5 min. Samples were then loaded on a 4%-12% NuPAGE gradient gel (Invitrogen #NP0233BOX). Gels were transferred onto a nitrocellulose membrane at 30V for 18 hrs. Membranes were blocked with 5% dry milk diluted in PBS containing 0.1% Tween 20 (PBS-T) for 1 hr at room temperature. Membranes were incubated in primary antibodies against DIAPH1 (Bethyl Laboratories A300-078A) and GAPDH (1:1000; Cell Signaling Technologies #2118L) diluted in 1X PBS-T containing 5% dry milk overnight at 4°C. Membranes were washed 3X for 5 min with 1X PBS-T and incubated in the secondary antibody IRdye 800 donkey anti-rabbit (LI-COR #926-32213) for 1 h at room temperature. Membranes were then washed 3X for 5 min in 1X PBS-T and imaged using the Odyssey CLx infrared scanner (LI-COR). Images were processed using the FIJI software (NIH) and protein expression levels were normalized to GAPDH.

### Light Microscopy and Image Processing

Laser scanning confocal imaging was conducted using a Nikon A1 Microscope equipped with 405, 488, 561, and 645 nm LASERs, Plan Apo 60X/1.4NA, and Plan Apo 25X/ 1.05 NA silicon (SIL) immersion objectives. Live-cell imaging was performed on a Nikon Ti2 inverted light microscope equipped with a Yokogawa CSU-X1 spinning disk head, equipped with 488 nm, 561 nm, and 647 nm excitation LASERs, a 405 nm photo-stimulation LASER directed by a Bruker mini-scanner to enable targeted photobleaching, a 100X Apo TIRF 100x/1.45 NA objective, and either a Hamamatsu Fusion BTor Photometrics Prime 95B sCMOS camera. Cells were maintained in a stage top incubator at 37°C with 5% CO2 (Tokai Hit). Super-resolution imaging was performed using a Nikon Structured Illumination Microscope (N-SIM) equipped with 405, 488, 561 and 640 nm LASERs, an SR Apo TIRF 100X/1.49 NA objective, and an Andor iXon Ultra DU-897 EMCCD camera. Images were reconstructed using Nikon Elements software. For imaging in all microscope modalities, imaging acquisition parameters were matched between samples during image acquisition. All images were denoised and deconvolved in Nikon Elements.

### Quantification and Statistical Analysis

All images were processed and analyzed using Nikon Elements software or FIJI software (https://fiji.sc/).

#### Analysis of length sum of filopodia overtime

A Filopodia Weka macro (Hebron et al., 2018) was used to classify the CDHR2TM channel of the motor domain movies into filopodia, cell body, or background through the Trainable WEKA Segmentation plugin (https://imagej.net/plugins/tws/). The subsequent generated probability maps for classified filpodia were then run through a length measurement macro, which auto threshold, convert to a mask, dilate, skeletonize and then run the Ridge Detection plugin (https://imagej.net/plugins/ridge-detection) to quantify the length of detected filopodia for the filopodia probability map for each time point.

#### Analysis of fraction change of filopodia number

The number of filopodia per cell at 0 min and 25 min either with or without rapalog treatment was counted, and the fraction change was quantified as the number of filopodia at 25 min over 0 min.

#### Measuring the length and rate of filopodia

Lengths of individual filopodia were measured in Fiji using the segmented line tool from the edge of the cell body to the end of the myosin-10 EGFP signal. Rate was calculated as the slope of the line fit via simple linear regression to the length over time.

#### Analysis of VASP tip enrichment in filopodia

VASP intensity along filopodia was measured using Nikon Elements software. A segmented line was drawn on filopodia from the tip of filopodia to the cell body, with either Myo10MD or Myo10FL at the tips. As all filopodia were different lengths or sometimes overlapped with other filopodia, measurements from the tip to ∼1.3 um towards the cell body were taken and normalized from 0 to 1, where 1 represented the filopodia tip. These normalized average intensity values were plotted and the line connects the average data points.

#### Quantification of the percent of cells exhibiting filopodia elongation

The percentage of cells exhibiting filopodia elongation in HeLa cells was calculated from HeLa cells expressing both the Myo10MD and either of the three membrane docking motifs. A HeLa cell was considered positive for filopodial elongation if the cell had at least four filopodia with the motor domain localized to the tip.

#### Length and either Myo10MD or Myo10-FL intensity measurements

ROIs of similar size were drawn over the tips of filopodia induced by either over expression of the Myo10MD-CDHR2TM system or full-length myosin-10 (Myo10-FL). Intensity measurements for either Myo10MD or Myo10-FL were taken, and the length of the associated filopodia was measured using the CDHR2TM channel in Fiji.

#### Analysis of filopodia length sum for Lat

*A washout and CellMask addition experiements.* For these experiments, the lengths of individual filopodia were taken manually for the indicated time points.

#### FRAP analysis

ROIs of similar area were drawn over the edge of HeLa cells expressing either of the three membrane binding motifs (CDHR2TM-FKBP, BTK-PH-FKBP, or FKBP-TH1) and bleached using a 405 nm LASER steered with a Bruker mini-scanner. Cells were imaged for 1 min prior to bleach and then imaged with no delay for 5 min to capture signal recovery dynamics. The first 2 min of recovery were used for analysis, and the first 50 sec of recovery are displayed on the graphs. All intensity values for each condition were normalized to the cell body intensity and background intensity as a ratio of (stimulated – background)/(cellbody – background). These FRAP intensities from multiple cells were then normalized from 0 to 1 and plotted together. Average values for each condition were fit using two-phase association curves. Immobile fractions were calculated as 1 minus the plateau of the curve.

#### Statistical analysis

Superplots were generated as described by (Lord et al., 2020). All experiments were completed in triplicate or more. Statistical significance was performed using the unpaired Student’s t-test for comparisons and the paired Student’s t-test for comparisons of the fraction change of the number of filopodia. All length filopodia sum over time statistical significance was performed using a two-way ANOVA followed by a Dunnett’s multiple comparisons test. All statistical analysis were computed using PRISM v.9.0. (GraphPad).

## SUPPLEMENTAL TABLE 1

**Supp. Table. S 1.**
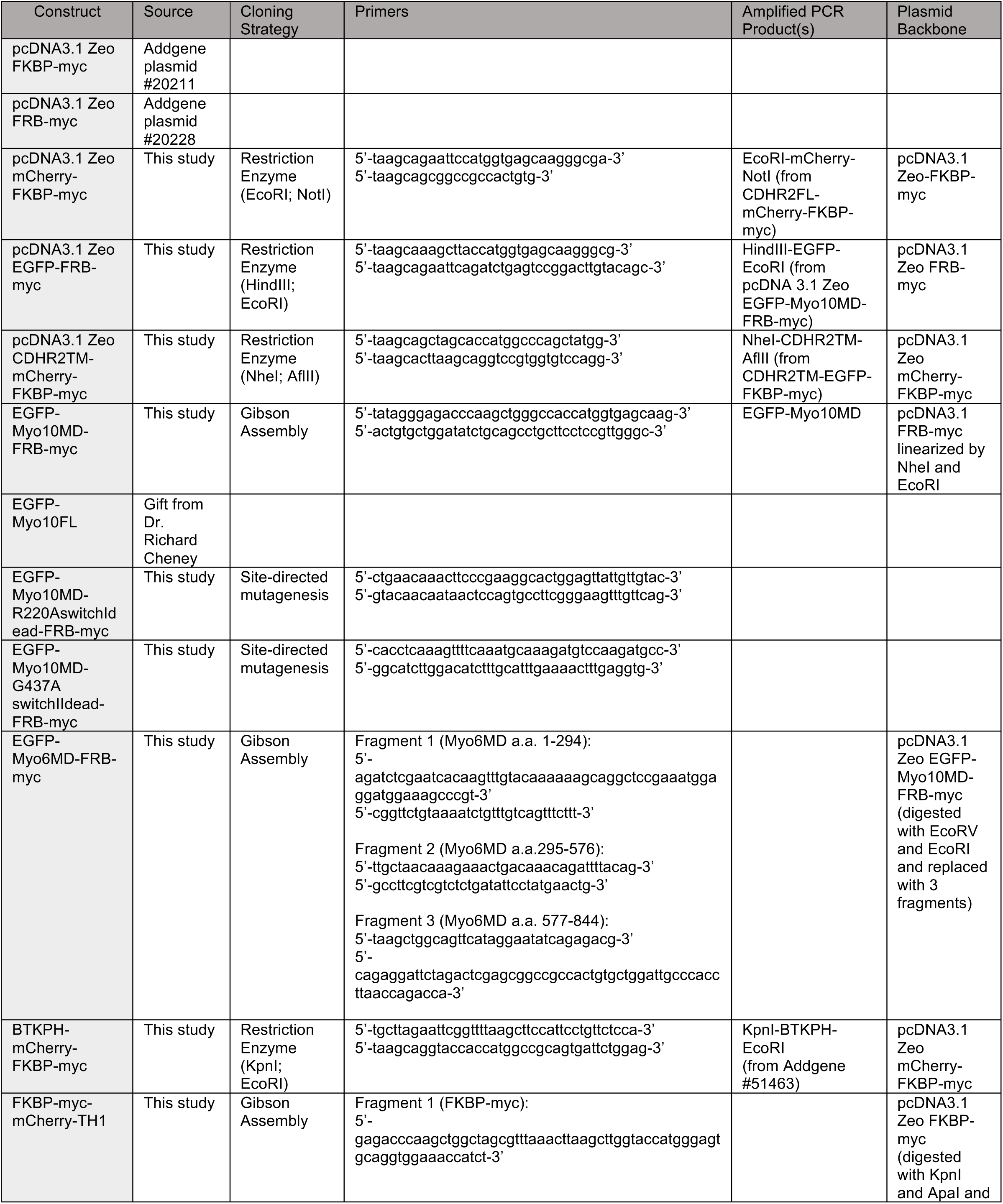
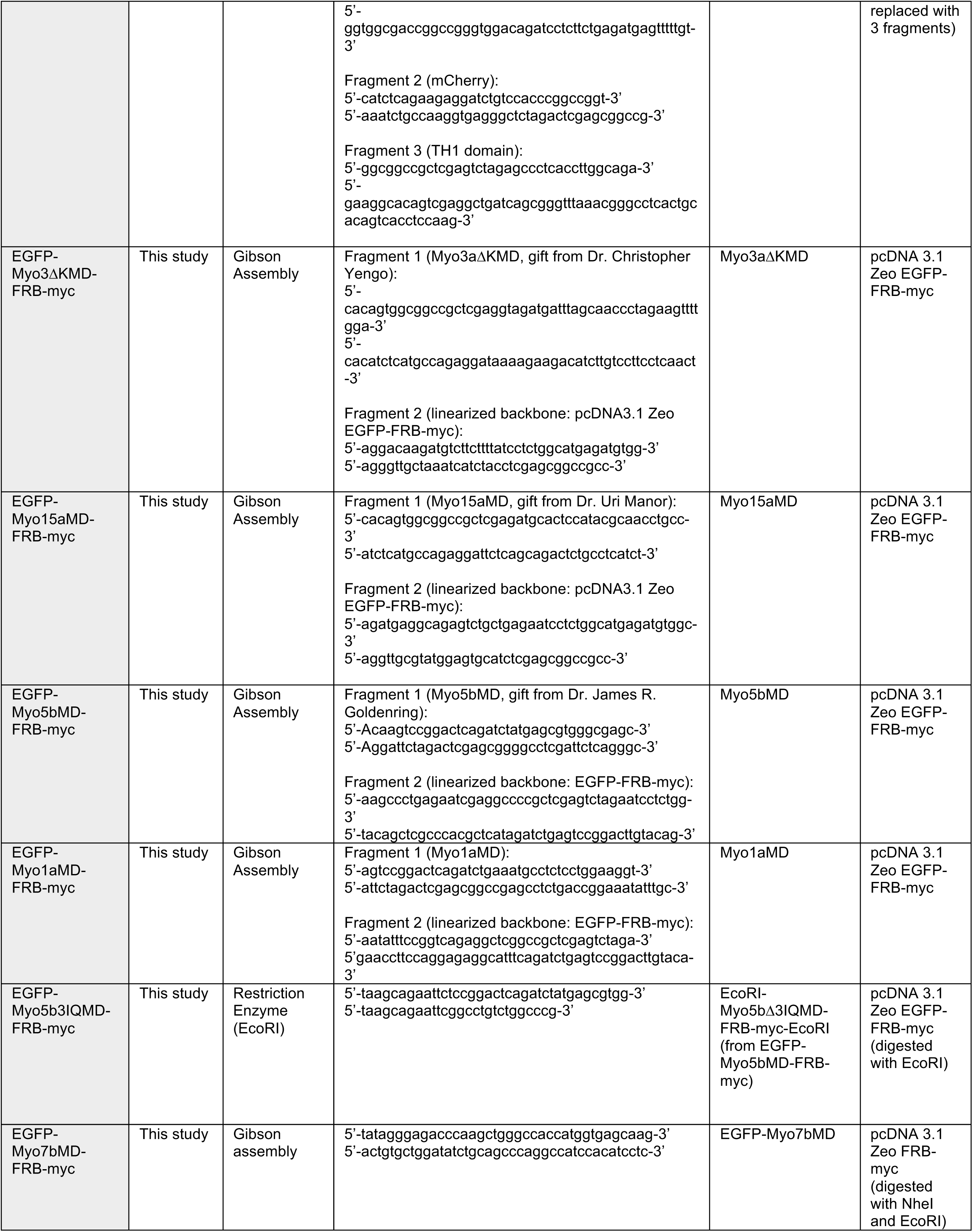

